# Hippocampal patterns and associative memory: Distinct intracranial EEG temporal encoding patterns support memory

**DOI:** 10.64898/2026.03.18.712716

**Authors:** Alice M. Xue, Shaw Hsu, Karen F. LaRocque, Omri Raccah, Alex Gonzalez, Josef Parvizi, Anthony D. Wagner

## Abstract

Episodic memory depends on neural representations encoded in the hippocampus. Experimental and computational evidence suggests that the hippocampus encodes pattern-separated representations that support later recall of episodic event elements. While extant data in humans predominantly focus on assaying the relationship between the similarity of spatial neural patterns at encoding and later memory performance, similarity of neural patterns in the temporal domain may also reveal encoding computations predictive of future memory. To examine how the similarity among temporal patterns of hippocampal activity during encoding relates to later episodic retrieval (associative cued recall and recognition memory), hippocampal activity was recorded from human participants (n=7) with implanted intracranial electrodes while they encoded arbitrary (A-B) paired-associates. Subsequent memory analyses first revealed that hippocampal high-frequency broadband power (HFB; 70-180Hz) was linked to a graded increase in memory strength; HFB power was greater during the encoding of pairs later correctly recalled relative to events later recognized and was lowest for events later forgotten. Second, and critically, subsequent memory analyses further revealed that more distinctive temporal patterns in the hippocampus during encoding — indexed by the similarity of the HFB timeseries elicited by a given event to that elicited by other events — were associated with superior subsequent memory performance. Finally, exploratory analyses revealed stimulus category effects on hippocampal HFB power during encoding and retrieval cuing. These results indicate that the temporal distinctiveness of hippocampal traces during encoding is important for subsequent retrieval of episodic event elements, consistent with theories that posit that pattern separation facilitates future remembering.

## Introduction

The ability to discriminate between similar past experiences is a fundamental function of episodic memory (e.g., remembering where your bike was parked today as opposed to yesterday). The hippocampus is theorized to support this function by forming a distinct neural representation for each experience. This ‘pattern separation’ process is thought to orthogonalize memories with similar input patterns to form distinct representations in the hippocampus. The orthogonalization of hippocampal representations is argued to minimize interference between memory traces, thus enabling later pattern completion that supports subsequent remembering of the details of a specific event (McClelland et al., 1995; Norman and O’Reilly, 2003; O’Reilly and McClelland, 1994; Treves and Rolls, 1994; Yassa and Stark, 2011).

To test this hypothesis, human electrophysiological and functional magnetic resonance imaging (fMRI) studies have examined the relationship between hippocampal activation patterns and memory behavior. Analyses of spatial activity patterns — across voxels (from fMRI) or channels (from intracranial electroencephalography; iEEG) — suggest that items (or events) elicit more distinct representations along the spatial dimension in the hippocampus than in other brain regions (Bakker et al., 2008; Berron et al., 2016; Jiang et al., 2020; Lacy et al., 2011; LaRocque et al., 2013). For example, using multivariate classification analyses over fMRI retrieval data, Chadwick, Hassabis, & Maguire (2011) observed that the spatial activity patterns associated with memory for four movies with overlapping content were more separable (i.e., decodable) in the hippocampus than in surrounding medial temporal cortical regions. Moreover and critically, studies quantifying pattern similarity during memory encoding indicate that later memory for an item/event is predicted by the extent to which its elicited spatial activity pattern in the hippocampus is distinct from the patterns elicited by other items/events (Ballard et al., 2019; Chanales et al., 2017; Favila et al., 2016; Hulbert and Norman, 2015; LaRocque et al., 2013). Taken together, extant studies provide evidence for the hippocampus’s role in encoding orthogonalized distributed spatial representations, which putatively reduces interference between similar memories and fosters later remembering.

Given that neural activity is organized both spatially and temporally as events unfold over time (Vaz et al., 2020; Yaffe et al., 2014), neural representations of episodic memories can also be characterized along the temporal dimension. Analyses of neural representations as arising from temporal sequences of spikes have been a cornerstone of the rodent hippocampal literature (Colgin et al., 2008). Yet, while prior studies in humans provide valuable insights about how the distinctiveness of hippocampal spatial patterns predict later remembering, they do not address whether distinctiveness in temporal coding within the hippocampus also relates to memory. One possibility is that hippocampal pattern separation mechanisms also occur over the temporal dimension, as suggested by emerging evidence that (a) temporal patterns of activity carry information about individual events (Pacheco Estefan et al., 2019; Staresina et al., 2016; Staudigl et al., 2015; Yaffe et al., 2014; Zhang et al., 2015) and (b) that at the single-cell level in isolated rodent hippocampal slices, namely in dentate gyrus and subfield CA3, similar temporal input patterns elicit more dissimilar temporal output patterns (Madar et al., 2019a, 2019b). Moreover, an iEEG study revealed that spatiotemporal patterns of activity recorded from human hippocampus exhibit evidence of pattern separation (Lohnas et al., 2018). However, because analyses in that study used patterns of activity combined over both spatial and temporal dimensions, it remains possible that the observed differences in spatiotemporal patterns were driven by pattern distinctiveness in space rather than in time or a combination of the two. It thus remains unclear whether the distinctiveness of temporal representations plays a role in hippocampal event encoding that supports later remembering.

Motivated by these observations, we leveraged the high temporal resolution of iEEG local field potential (LFP) recordings of high-frequency broadband power (HFB; 70-180Hz), which is thought to reflect local population activity (Buzsáki et al., 2012; Manning et al., 2009; Miller et al., 2014; Mukamel et al., 2005). We recorded hippocampal LFPs from seven epileptic patients while they were engaged in a blocked study-test associative memory paradigm to examine the link between distinctiveness of elicited temporal neural patterns at encoding and later memory performance (Fig. 1A; see Supplementary Material, Figure S1, Table S1, and *Materials and Methods* for participant demographics and analytic approach on electrodes with clinical pathology). During the encoding phase, participants studied arbitrary (A-B) pairs of items that were presented sequentially (items were drawn from one of four stimulus categories; see *Materials and Methods*). During the retrieval phase, participants encountered studied items (the As) and novel items and (a) made old/new recognition memory judgments and (b) engaged in cued recall of studied associates (the Bs). Critically, to isolate the temporal component of the hippocampal representation, distinctiveness was calculated from similarities of temporal patterns at individual recording channels. To the extent that hippocampal pattern separation is evident in temporally unfolding neural patterns, we hypothesized that events that elicited more distinct temporal patterns of activity at encoding would be more likely to be later remembered. We also examined hippocampal univariate HFB power via a priori analyses of effects of subsequent memory (Greenberg et al., 2015; Long et al., 2014; Staresina et al., 2016) and exploratory analyses of effects of stimulus category.

**Figure 1.**
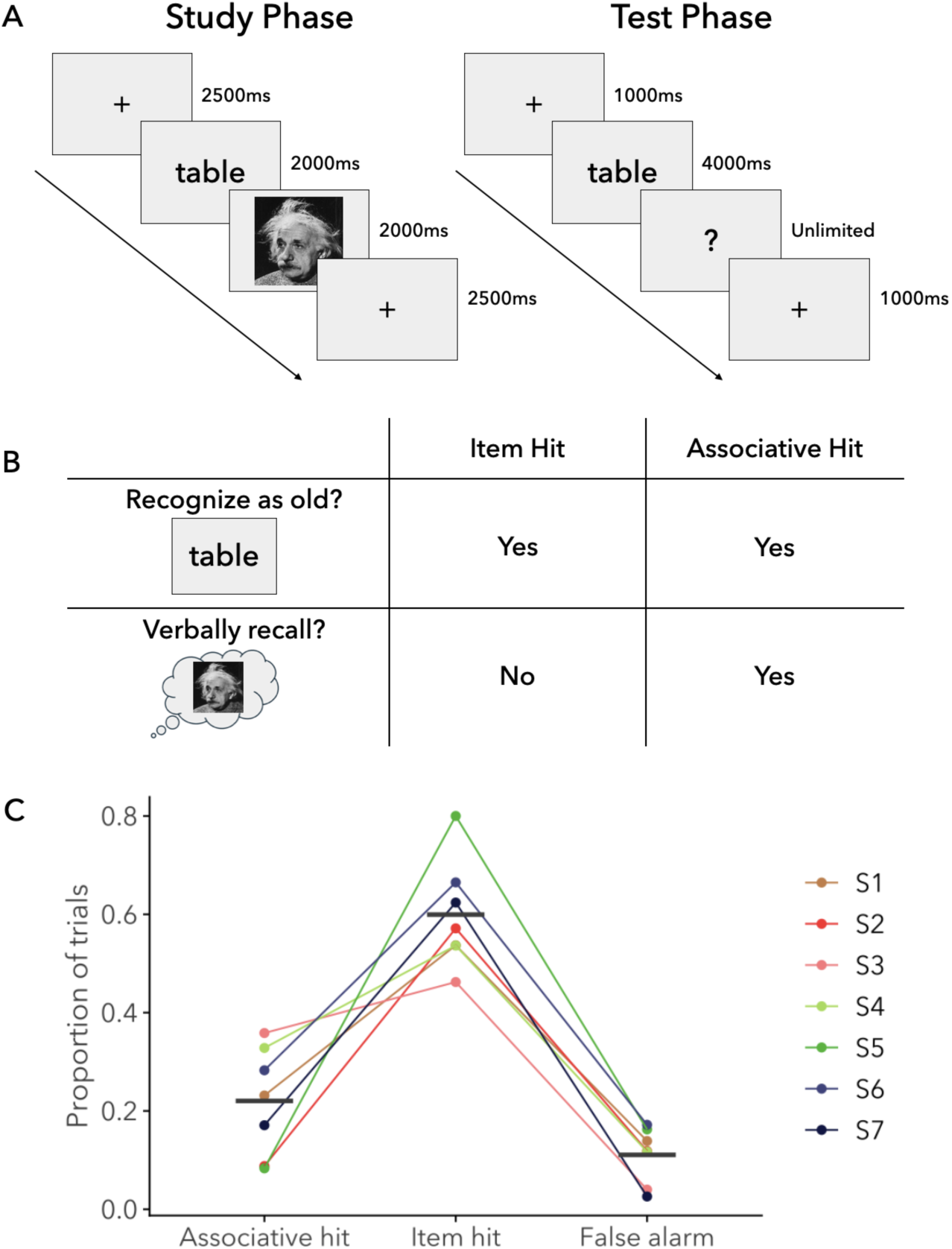
Experimental design and behavioral results. (A) Study phase: In each block, 12 pairs of items were sequentially presented at the center of the screen, and participants were instructed to memorize the item A-item B pairings. Test Phase: In each block, 20 cues (12 old and 8 new) were presented individually, and participants were instructed to indicate whether each cue was old/new and to then verbally recall the associated item if they recognized the cue as old. (B) Definitions of item hit and associative hit categories. (C) Proportions of behavioral outcomes at retrieval. Each point denotes an individual participant and black line segments denote the means. Associative hit: proportion of old cues for which participant could recall the associate; Item hit: proportion of old cues recognized as studied, but without recall of the associate; False alarm: proportion of new cues falsely recognized as studied.

## Materials and Methods

### Participants

Seven human epileptic patients (Table 1) participated in the experiment. Patients were implanted with depth electrodes as part of a presurgical evaluation to localize their seizure onset zone; locations of implantations were determined solely based on clinical considerations, and participation in the experiment did not carry any additional risk to the patients. All patients gave informed written consent in line with procedures approved by the Stanford Institutional Review Board.

**Table 1.** Participant demographics and subsequent memory performance. Recognition memory was computed as d’ and cued recall as associative hit rate and associative error. rate Full-4 IQ was obtained from the Wechsler Abbreviated Scale of Intelligence (WASI-II) test for 6 of the 7 participants. Right/left and anterior/posterior hippocampus electrode location for each participant are shown.

| Participant | Age | Sex | Number<br>of blocks | d' | Assoc. hit rate | Assoc. error rate | Full-4 IQ | R Ant. | L Ant. | R Pos. | L Pos. |
| --- | --- | --- | --- | --- | --- | --- | --- | --- | --- | --- | --- |
| S1 | 43 | F | 9 | 1.789 | 0.231 | 0.014 | 86 | 2 | 0 | 0 | 0 |
| S2 | 33 | F | 10 | 1.564 | 0.088 | 0 | 67 | 2 | 2 | 0 | 0 |
| S3 | 41 | M | 10 | 2.591 | 0.358 | 0 | 121 | 4 | 2 | 0 | 2 |
| S4 | 43 | M | 16 | 2.267 | 0.328 | 0.023 | 99 | 3 | 0 | 0 | 0 |
| S5 | 26 | M | 10 | 2.143 | 0.083 | 0 | 78 | 2 | 0 | 0 | 0 |
| S6 | 44 | F | 16 | 2.538 | 0.283 | 0.023 | – | 2 | 0 | 0 | 0 |
| S7 | 19 | F | 10 | 2.666 | 0.171 | 0 | 91 | 0 | 2 | 0 | 7 |
| Mean(SD) |  |  |  | 2.223(0.421) | 0.220(0.111) | 0.009(0.011) | 90.3(18.6) |  |  |  |  |

### Neuropsychological Testing

Patients were administered the California Verbal Learning Test-II (CVLT-II) and the Wechsler Abbreviated Scale of Intelligence (WASI-II).

### Experimental Paradigm and Task Design

Participants performed a paired-associate paradigm that consisted of multiple blocks of study (encoding) and test (retrieval) (Fig. 1A and Table 1). During the study phase for each block, participants viewed 12 pairs of items. Each arbitrary pair consisted of two unique items (A and B) that were presented sequentially for 2 s each, followed by an intertrial interval (fixation cross; 2.5 s). Participants were explicitly told that their memory for each pair would be tested in the next round of the experiment.

Between the study and test phases for each block, participants completed an arrow detection distractor task for 15 s. During the arrow detection distractor task, participants were presented with a series of arrows, each pointing to the left or right with equal probability. Participants were instructed to respond as quickly and as accurately as possible, pressing one of two buttons that corresponded to the direction of the arrow presented on the screen. Each self-paced trial was followed by an intertrial interval of 0.5 s. The distractor task terminated after 15 s. During the test phase for each block, participants were tested on their memory for the items presented during the study phase for that specific block. Participants were shown 20 cue items, which consisted of a mixture of all 12 A items from the study phase and an additional 8 novel foils; B items from the study phase were never presented as the cue item in the test phase. For each cue item, participants could respond (1) “new,” (2) “old, remember nothing about its pair,” or (3) “old, remember something about its pair” by pressing one of three buttons. Participants were instructed to respond within the 4 s while the cue was present on screen; presentation of each cue terminated following a participant’s recognition memory response or 4 s, whichever came sooner. If participants responded with (3) or if they failed to respond within 4 s, a question-mark was presented and the experimenter verbally probed and recorded the participant’s memory for the paired associate (i.e., the B item). Once the verbal probe period concluded, the experimenter manually resumed the experiment. Each test trial was followed by a 1-s intertrial interval (fixation).

Prior to beginning the experiment, participants practiced the task on a shortened list that consisted of 4 study pairs and 6 test cues (4 old + 2 novel foils). Participants performed the practice and experimental tasks on an Apple laptop. Stimulus presentation and participant response recording was conducted using PsychToolBox in MATLAB. Participants performed a maximum of 16 study-test blocks in the experiment, with the number of blocks performed determined by clinically available recording time (Table 1).

### Stimuli

All items belonged to one of four categories: pictures of famous faces, pictures of famous buildings, numbers (2 digits in blocks 1-11; 3 digits in blocks 12-16), and words (nouns). All stimuli were presented centrally on a gray background in grayscale.

During each study phase, items from the same category never occurred in the same pair. For each block, the 12 studied pairs consisted of all 12 possible pair-combinations of the 4 categories excluding same category pairs. This resulted in each category being the A term in 3 pairs and the B term in 3 pairs, thus allowing exploratory analyses of stimulus category coding in hippocampal encoding and retrieval activity (Supplementary Material and Fig. S6). All stimuli used for each participant were unique within each block and across all blocks of the experiment (i.e., items never repeated). The stimuli used for each participant were randomly drawn from an overall pool of stimuli, with the assignment (pairs; and old vs novel at test) and presentation order of stimuli randomized across participants.

### Data Recording

Data were recorded using implanted depth electrodes using a multi-channel recording system (Tucker-Davis Technologies). The depth electrodes were cylindrical, 1.3 mm in diameter with 1.2 mm^2^ of exposed recording area and an inter-electrode distance of 5 mm. Data were recorded at a sampling rates of 3051.8 Hz (participants S1, S2, and S3; sampling rate was subsequently upsampled to 1000Hz to match the other participants) or 1000 Hz (participants S4, S5, S6, and S7). Data were analyzed offline using custom software in MATLAB and R.

### Preprocessing

Data from each electrode were linearly detrended, bandpass filtered between 1 Hz and 180 Hz, notch filtered at 60 Hz and higher harmonics to remove line noise. Data were then re-referenced to the common average signal from the set of all electrodes located in white matter (mean= 16; range= 5-30). To obtain HFB (70-180 Hz), a Morlet wavelet transform (wavelet number: 6) was applied to the signal at 5 Hz intervals to obtain narrow band signals. Each narrow band signal was normalized by dividing by its own mean amplitude. The normalized signals were averaged and multiplied by the mean amplitude across all frequencies to bring the signal back into volts. The amplitude was then squared to obtain power and natural log transformed. The above normalization method aimed to correct for the undesired 1/f decay observed in the EEG power spectrum that results in dominance of lower frequency components (Buzsáki and Mizuseki, 2014; Norman et al., 2017; Preston et al., 2025). Photodiode markers were used to epoch the signal into individual trial epochs. Each epoch was normalized by subtracting off the mean log HFB in the 200 ms preceding each epoch.

Electrodes were selected for analyses if they were unambiguously located in the hippocampus and if they were not contaminated with machine noise. Rather than discarding electrodes based on clinical pathology, we used a stringent artifact detection procedure to eliminate trials showing abnormal electrophysiological activity. Artifact detection was performed on the signal after each step of preprocessing. Separately for the raw data, filtered data, re-referenced data, and log HFB, time points were marked as artifacts if they exceeded five standard deviations from the signal mean or if their temporal derivative (change in magnitude from one time point to the next) exceeded five standard deviations from the mean temporal derivative. For each artifact time point, an artifact window that contained all time points within 200 ms of the artifact time point was created. All trial epochs that contained any temporal overlap with any artifact window were excluded from analyses. Trial epochs were excluded separately for each electrode. Electrode localization and selection were guided by post-surgical CT images of electrodes that were aligned with structural MRI of the brain for each participant. Alignment was performed using a mutual-information algorithm implemented in SPM (http://www.fil.ion.ucl.ac.uk/spm/).

### Behavioral Analysis Methods

The memory outcomes for each test trial were initially categorized into eight memory categories, which were further reduced to five categories for the analyses presented in the results section. The initial behavioral outcomes were: (1) miss: responded to a studied cue with “new”; (2) item hit: responded to a studied cue with “old, remember nothing about its pair”; (3) item hit + associative category miss: responded to a studied cue with “old, remember something about its pair” but failed to correctly identify the category of the pair during verbal probe; (4) item hit + associative category hit: responded to a studied cue with “old, remember something about its pair” and correctly identified the category of the pair but was unable to correctly generate at least one qualitative descriptor of the associate during verbal probe; (5) associative hit: responded to a studied cue with “old, remember something about its pair,” correctly identified the category of the pair, and correctly generated at least one qualitative descriptor of the associate during verbal probe; (6) correct rejection: responded to a novel cue with “new”; (7) item false alarm: responded to a novel cue with “old, remember nothing about its pair”; and (8) associative false alarm: responded to a novel cue with “old, remember something about its pair.” Due to low trial counts for some categories, categories (2), (3), (4) were combined into the “item hit” category and categories (7) and (8) were combined into the “false alarm” category for the analyses presented in the Results section.

Sensitivity index d’ was used to evaluate recognition memory performance and was calculated as d’ = Z(total hit rate) – Z(false alarm rate), where the total hit rate was calculated as the sum of the number of item hits and associative hits divided by the total number of studied items and the false alarm rate was calculated as the number of false alarms divided by the total number of lure items. Z is the inverse of the cumulative distribution function of the Gaussian distribution. Associative hit rate was used to evaluate cued-recall performance and was calculated as the number of associative hits divided by the total number of studied items. Associative recall error rate was calculated to be identical to the associative false alarm rate.

For models that involved multiple measurements per participant, Bayesian linear mixed-effects models that treated participants as a random effect were used and implemented using the brms package in R (Bürkner, 2017). To assess the importance of predictors of interest, Bayesian leave-one-out cross-validation of models including and excluding the predictor of interest was performed to test the generalizability of the models on out-of-sample data. Predictive performance of each model was assessed using Expected Log Predictive Density (ELPD) (Vehtari et al., 2017). In the main text, ELPD for the better performing model is reported alongside the difference in ELPD between the two models and the associated standard error. Analyses involving only a single measurement per participant were performed using standard Bayesian linear models with a sole intercept term. Group measurements were considered credibly different from zero if the 95% highest density interval (HDI) excluded zero.

### HFB Analyses

All Bayesian linear mixed-effect models used in the temporal pattern similarity and univariate analyses included nested participant and electrode random intercepts. For study phase item A-period pattern similarity analyses, the models included item A category and item A mean univariate HFB amplitude as control variables. For study phase item B-period pattern similarity analyses, the models included both item A and item B categories and both item A and item B mean univariate HFB amplitudes as control variables. For all univariate HFB analyses, HFB was first divided into 100-ms time bins separately for each electrode. For study phase item A univariate analyses, the models included time-bin, item A category, and subsequent memory outcome as predictors and controls. For study phase item B univariate analyses, the models included time-bin, both item A and item B categories, and subsequent memory outcome as predictors and controls. For test phase univariate analyses on the effects of item categories, the lure trials were excluded, and the model included time-bin, item A and item B categories, and subsequent memory outcome as predictors and controls. For test phase univariate analyses on the effects of memory outcome, all trials were included, and time-bin and item A category were included as control variables. Smoothing of the timeseries of univariate HFB amplitudes was performed only for visualization using thin plate spline regression.

The importance of predictors of interest was assessed using Bayesian leave-one-out cross-validation, as described above (see *Behavioral Analysis Methods*). Comparisons between conditions of interest were performed by computing estimated marginal contrasts using the emmeans package (“Estimated Marginal Means, aka Least-Squares Means,” n.d.). Differences between conditions were considered credible if the 95% HDI excluded zero. The posterior median and 95% HDIs for the estimated marginal contrasts are reported in the text.

## Results

### Subsequent Memory Performance

Recognition memory and cued-recall performance at test focused on three memory outcomes (Fig. 1; Fig. S2): associative hit rate/cued-recall probability (proportion of studied cues/A items correctly recognized as ‘old’ and for which the paired associates/B items were recalled); item hit rate (proportion of studied cues correctly recognized as ‘old’ but for which the paired associates were not recalled), and false alarm rate (FA, proportion of novel cues incorrectly recognized as ‘old’). By definition, item hits reflected a mixture of decisions based on recollection of other event features (i.e., features other than specifics about the B item) and on item familiarity.

To quantify participants’ subsequent memory performance, we computed the sensitivity index (d’) from recognition memory judgments on test cues (A items) and the probability of cued recall of the associates (B items) via associative hit rate (Table 1; Fig. 1C). Analyses indicated that recognition memory (d’) was above chance (M = 2.22 [1.80, 2.64]), and that associative recall rate was above zero (M = 0.22 [0.11, 0.33]) with very low associative error rates (M = 0.009, SD = 0.011). Importantly, all participants had recognition memory d’s above 1.5, indicating strong discriminability in memory performance. Model comparison of linear mixed-effects models examining memory performance revealed that including study-test block as a predictor did not meaningfully improve predictive performance for recognition memory (d’) (Expected Log Predictive Density (ELPD) for model without block = −73.40, ΔELPD = −1.33± 0.84), indicating that participants performed consistently across the experiment. We similarly examined whether recognition memory and cued recall varied as a function of encoding serial position within a study-test block (Fig. S3). Model comparison of mixed-effects logistic regression models on miss rate (the complement of recognition of studied cues) and on associative hit rate, with or without encoding serial position entered as a categorical predictor, revealed that neither recognition memory (ELPD for model without serial position = −391.52, ΔELPD = −8.77 ± −2.76) nor cued recall (ELPD for model without serial position = −484.83, ΔELPD = −6.06 ± 3.47) varied meaningfully with serial position. Additional exploratory analyses examined the effects of stimulus category on memory performance (see Supplementary Material and Fig. S4, S5).

We note that the well-above chance recognition d’s and low associative recall error rates suggest that participants with lower associative error rates (S2 and S5, in particular) may have adopted high response thresholds rather than having poor memory. Consistent with this interpretation, we note that 6 of 7 participants completed immediate and 20-minute delayed recall from the California Verbal Learning Test-II (CVLT-II), and the ratio of delayed to immediate recall was high for all participants ranging from 1.07 to 0.64 (with retention for S2 = 0.85; S5 = 0.80). At the same time, we note that Wechsler Abbreviated Scale of Intelligence (WASI-II) IQ scores were in the lower range for S2 and S5 (Table 1); for this reason, we confirmed that the main temporal pattern similarity findings reported below hold when the analyses were computed while excluding the data from these two participants (see Supplementary Material).

### Hippocampal HFB Power and Memory Performance

At the neural level, we first examined the relationships between univariate hippocampal HFB power and memory behavior. A priori analyses focused on how hippocampal HFB power at encoding varies with subsequent memory (Brewer et al., 1998; Davachi et al., 2003; Long et al., 2014; Staresina et al., 2016; Wagner et al., 1998); for completeness we also report HFB power at retrieval. For the study phase, the item A and item B trial periods were analyzed separately (each using 2-s post-item onset). The test phase data were analyzed both as a function of cue onset (cue-locked; 1-s post-cue) and participant response (response-locked; 1-s preceding response). Separate linear mixed-effects models linking HFB and memory outcomes with different sets of predictors and control variables were constructed with nested participant/electrode random effects. Bayesian leave-one-out cross-validation based model comparison was used to evaluate predictors of interest. Estimated marginal contrasts for predictors of interest for each model and HFB time courses are shown in Fig. 2.

**Figure 2.**
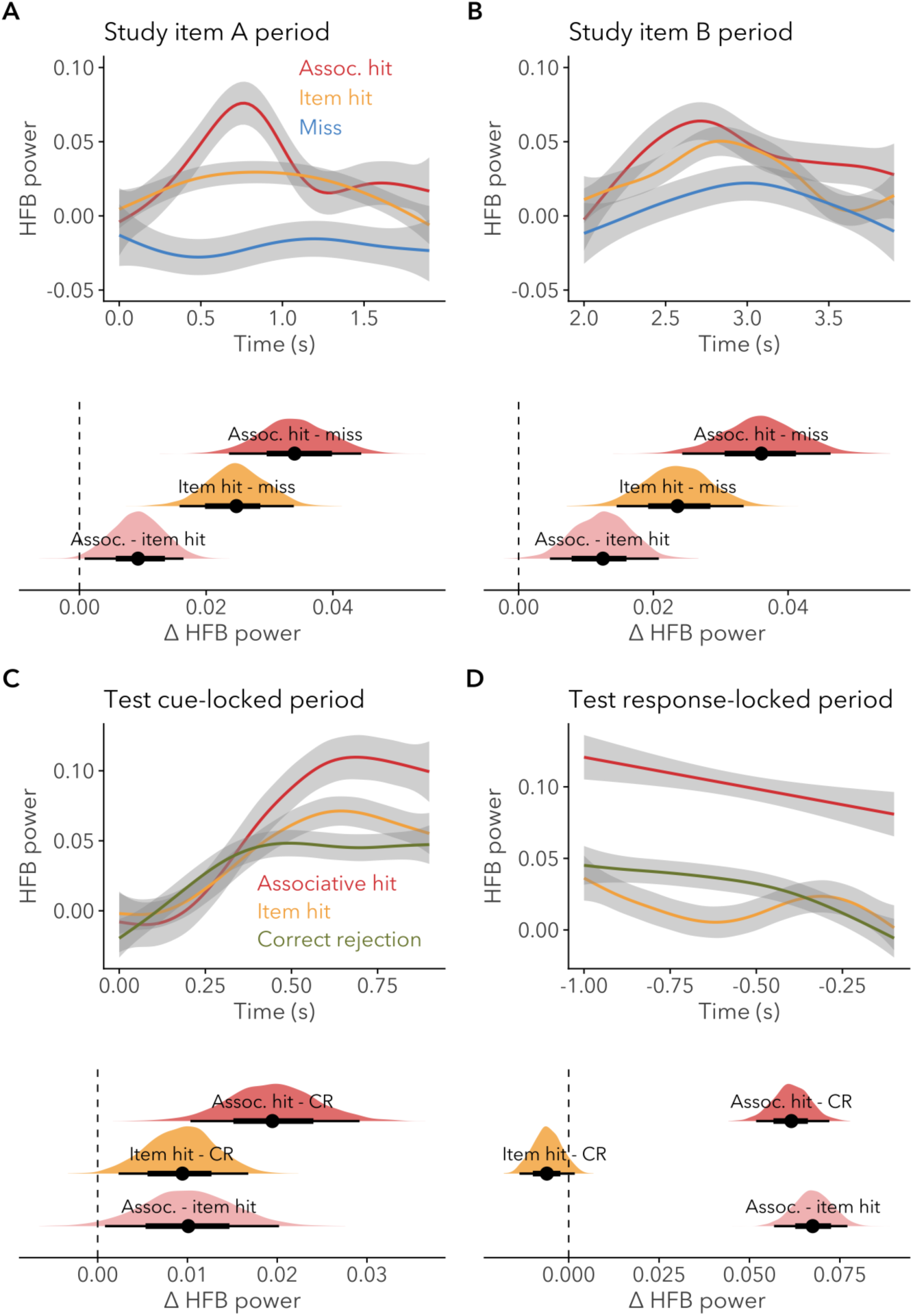
Univariate HFB timeseries plots (top) and posterior probability distributions for estimated marginal contrasts for HFB between memory outcomes (bottom) during the study phase (A, B) and test phase (C, D). Timeseries were smoothed for visualization purposes. Error bars on the timeseries represent bootstrapped 95% confidence intervals. Plots of estimated marginal contrasts show posterior probability distributions in the shaded areas; points indicate posterior medians and horizontal lines indicate the 66% and 95% HDIs.

For the study phase (Fig. 2A, B), inclusion of subsequent memory as a predictor meaningfully improved predictive performance for both the item A (ELPD = −19332.07, ΔELPD = −18.41 ± 6.37) and item B (ELPD = −21095.94, ΔELPD = −18.09 ± 6.27) encoding periods. Follow-up analyses of estimated marginal contrasts showed that (a) HFB power for subsequent associative hits was higher than both subsequent item hits and misses during encoding of item A (Δ = 0.009 [0.001, 0.017]; Δ = 0.034 [0.024, 0.044]) and during encoding of item B (Δ = 0.013 [0.004, 0.020]; Δ = 0.036 [0.024, 0.046]); and (b) HFB power for item hits was higher than misses during encoding of item A (Δ = 0.025 [0.016, 0.034]) and during encoding of Item B (Δ = 0.024 [0.014, 0.033]). For the test phase (Fig. 2C, D), inclusion of memory (i.e., retrieval success) as a predictor improved predictive performance for both cue-locked (ELPD = −15787.97, ΔELPD = −8.48 ± 4.98) and response-locked periods (ELPD = −19910.82, ΔELPD = −102.76 ± 14.61). Follow-up analyses showed that (a) HFB power for associative hits was higher than CRs during the cue-locked (Δ = 0.019 [0.011, 0.030]) and response-locked periods (Δ = 0.062 [0.052, 0.072]); (b) HFB power for associative hits was higher than item hits for the response-locked period (Δ = 0.067 [0.057, 0.078]), but not the cue-locked period (Δ = 0.010 [0.000, 0.020]); and (c) HFB power for item hits was higher than CRs during the cue-locked period (Δ = 0.009 [0.002, 0.017]) but not the response-locked period (Δ = −0.006 [−0.014, 0.002]). Exploratory analyses examined the effects of stimulus category on hippocampal univariate HFB (see Supplementary Material and Fig. S6, S7).

### Hippocampal HFB Temporal Pattern Similarity and Subsequent Memory

Our central analyses used the subsequent memory approach (Paller and Wagner, 2002) to examine the relationship between the distinctiveness of hippocampal HFB temporal patterns at encoding and later differences in memory performance at test. For each electrode, the HFB temporal pattern for each item was defined as a vector of HFB values for that item; for each 2-s item presentation at study, the time course of HFB activity was quantified using 10-ms non-overlapping time bins, resulting in a vector of 200 elements for each item’s presentation. Critically, HFB temporal patterns were computed separately for each electrode, ensuring independence from spatial pattern similarity.

Using HFB temporal patterns, we generated the primary measure of interest — the *within-block temporal pattern similarity (wTPS)* index — for each study item to quantify the degree of similarity between the temporal pattern of that study item and the temporal patterns of all other items presented in the same block. More specifically, the wTPS for each study item was calculated as follows: (1) pattern similarity was computed as the Pearson correlation between the target item temporal pattern vector and the temporal pattern vector of each of the other study items in the same study block (A items were correlated with the other A items; B items with the other B items); (2) the wTPS score for the study item was calculated as the mean of all the correlation scores in step (1); and (3) each study item’s wTPS score was subsequently z-scored with respect to all other wTPS scores from the same block. Thus, wTPS for a study item reflects the degree to which that item’s evoked HFB timeseries is similar to those of all the other items in the same block. The wTPS scores were calculated separately for item A and item B presentations at encoding.

To examine the relationship between wTPS at study and subsequent memory outcome, linear mixed-effects models were used to regress wTPS on subsequent memory outcome (associative hit, item hit, miss), with nested participant-electrode random effects. Stimulus category as well as univariate HFB power were included in all models as controlling variables, regardless of whether these predictors were found to meaningfully improve predictive performance in the preceding univariate analyses. Importantly, consistent with our hypotheses, the model with item A wTPS as the outcome and subsequent memory as a predictor showed higher predictive performance (Fig. 3A; ELPD = −3406.66), though the difference in predictive performance was small (ΔELPD = −3.26 ± 3.14). Estimated marginal contrasts showed that item A wTPS for subsequent misses was greater than that for subsequent associative hits (Δ = 0.137 [0.022, 0.257]) and item hits (Δ = 0.173 [0.069, 0.275]), which did not differ from each other (associative hit – item hit = 0.036 [−0.044, 0.128]). Moreover, mean item A wTPS for subsequent associative hits was lower than that for subsequent misses in 7 of 7 participants, and mean wTPS for subsequent item hits was lower than that for misses in 6 of 7 participants. By contrast, separate models of item B wTPS revealed that predictive performance was higher when memory outcome was excluded as a predictor (Fig. S8; ELPD = −3414.07, ΔELPD = −1.70 ± 0.87); this indicates that pattern distinctiveness during item B presentation did not differ meaningfully between subsequent memory outcomes. Collectively, these results suggest that increased distinctiveness of hippocampal HFB temporal patterns for item A is correlated with superior subsequent memory performance.

**Figure 3.**
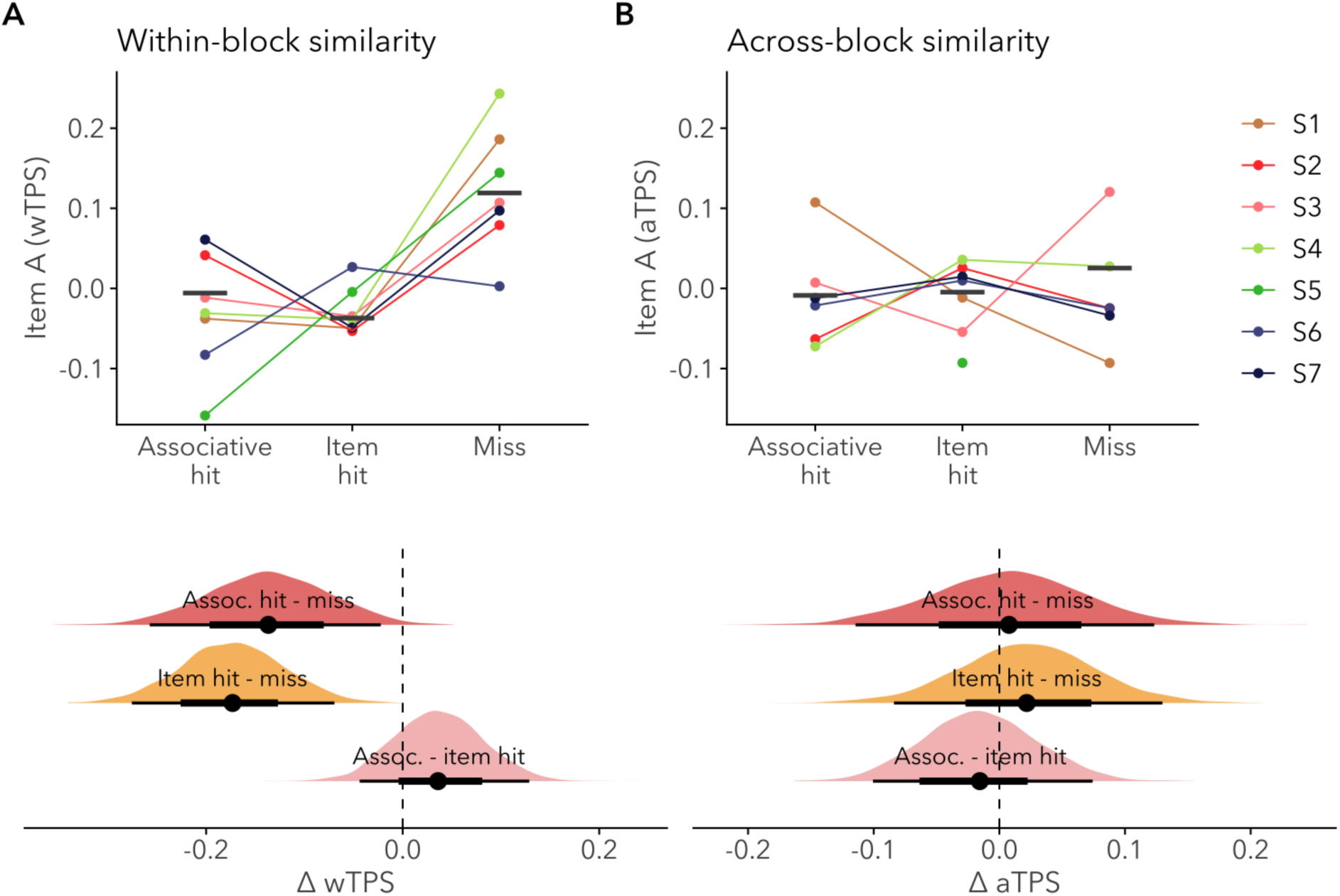
Study phase item A (A) within-block temporal pattern similarity (wTPS) and (B) across-block temporal pattern similarity (aTPS). Each point denotes an individual participant and black horizontal line segments denote weighted group means. Note: The data points for participant S5 in the Associative Hit and Miss columns are not visible in B due to low trial numbers causing outlier data points; see Fig. S9 for the complete plot and Supplementary Material for supplementary analyses. Estimated marginal contrasts between subsequent memory outcomes for within-block and across-block similarity are shown at the bottom of (A) and (B), respectively. Plots of estimated marginal contrasts show posterior probability distributions in the shaded areas; points indicate posterior medians and horizontal lines indicate the 66% and 95% HDIs.

One potential explanation for the observed relationship between lower wTPS at study and subsequent memory performance is that items that elicited more distinct temporal patterns at study suffer from less interference at later memory retrieval and thus are better remembered. However, an alternative to this temporal pattern separation account is that items that were more strongly encoded during study happened to elicit temporal patterns that contained some memory-independent temporal signature (e.g., a transient burst in HFB post-stimulus onset). This would cause the strongly encoded items to have more distinct temporal patterns from ‘normative’ (i.e. more typical) temporal responses, and this difference would also be reflected as a negative correlation between wTPS and subsequent memory performance. These two alternative accounts, however, make differential predictions about a study item’s distinctiveness when compared to study items in other blocks. The temporal pattern separation account would predict that a study item’s temporal pattern distinctiveness to items in future blocks should not relate to later remembering for that item because items from future blocks cannot cause interference with the current item in a blocked study-test design. The memory-independent temporal signature account would predict that the relationship between temporal distinctiveness and later memory does not depend on which block the temporal similarity was computed against, since the memory-independent temporal signature should be evident when computed against other study blocks.

To differentiate between these accounts, we generated an *across-block temporal pattern similarity* (aTPS) index to quantify the degree of temporal pattern similarity between a target study item and all items in the subsequent study block. The aTPS is computed in the same way as wTPS, with the exception that all correlations are calculated with respect to items in the subsequent study block relative to the target item. Thus, aTPS is a measure of how distinctive a study item’s temporal pattern is from the temporal patterns elicited from the subsequent block. To examine the relationship between aTPS at study and subsequent memory outcome, linear mixed-effects models were used to predict aTPS as a function of memory outcome (associative hit, item hit, miss), with nested participant-electrode random effects. Again, stimulus category and univariate HFB amplitude were controlled for in the models. Importantly, model comparison showed that item A aTPS was better modeled when subsequent memory outcome was excluded as a predictor (Fig. 3B; ELDP = −3414.37, ΔELPD = −1.74 ± 0.48); this suggests that aTPS during item A presentation did not differ meaningfully between subsequent memory outcomes. Moreover, mean item A aTPS for subsequent associative hits was lower than that for subsequent misses in only 3 of 7 participants, and mean aTPS for subsequent item hits was lower than that for misses in only 2 of 7 participants. While power to detect interactions is modest due to the sample size (which, while reasonably large for an iEEG study, was constrained due to patient availability), additional models showed that inclusion of the similarity type (wTPS vs aTPS) × memory outcome interaction did not improve predictive performance (ELPD for model without the interaction = −6820.96, ΔELPD = −0.25 ± 1.78) for the associative hit and miss contrasts and item hit and miss contrasts (Δ = −0.089 [−0.248, 0.059]; Δ = −0.123 [−0.262, 0.009]). To address potential concerns about the lack of statistical power due to low trial counts and to ensure that the results presented here are not driven by the specific memory outcome categorizations used, we recomputed the pattern similarity analyses both after combining associative hits and item hits into a single hit memory category and after categorizing memory outcomes in an alternative method and showed that the results hold in both cases (see Supplementary Material, Fig S1B, S10-12). Acknowledging that some caution is warranted given the null interactions, these results nonetheless lend support for the temporal pattern separation account, wherein greater hippocampal pattern distinctiveness is associated with superior subsequent cued recall and item recognition because events with lower wTPS putatively suffer from less interference from other events in the same study-test context.

## Discussion

Leveraging human iEEG recordings and temporal pattern similarity analyses, the present experiment revealed a relationship between the distinctiveness of hippocampal temporal patterns at encoding and subsequent memory performance. More specifically, relative to events later forgotten, events later remembered were associated with more distinct hippocampal temporal patterns at encoding. Comparisons of wTPS and aTPS offer further evidence that increased distinctiveness of hippocampal temporal patterns at encoding reduces interference between event representations, thus facilitating later retrieval. Together, these results complement computational and experimental research that implicates the hippocampus in creating distinctive (pattern-separated) memory representations to minimize interference (Knierim and Neunuebel, 2016; Kumaran et al., 2016; Yassa and Stark, 2011). The present findings substantively extend current understanding of hippocampal processes, providing evidence in support of (a) pattern separation in the human hippocampus, which has recently been debated (Quian Quiroga, 2020; Rolls, 2021; Suthana et al., 2021) and (b) the novel, critical insight that the relationship between pattern distinctiveness and memory operates not only over spatial patterns of neural responses (Bakker et al., 2008; Berron et al., 2016; Chanales et al., 2017; Favila et al., 2016; Hulbert and Norman, 2015; Lacy et al., 2011; LaRocque et al., 2013), but also over representations constructed from temporal patterns of neural activity (Madar et al., 2019a, 2019b).

How might populations of neurons in the hippocampus give rise to temporal patterns with varying degrees of distinctiveness that would be related to subsequent memory behavior? One possibility is that, even if the group of neurons that are activated by two different stimuli are the same ensemble, changes in the temporal patterns of activity can arise from changes in the firing order of the same set of neurons. If each neuron fires with a distinct signature, then the temporal representation elicited by each stimulus can be orthogonalized by changing the order in which the neurons fire. Recent data from humans engaged in a memory task showed that groups of hippocampal neurons rearranged their temporal firing order in response to presentations of different stimuli (Vaz et al., 2020). This provides support for interpreting the present changes in temporal patterns as changes, at least in part, in the underlying firing sequence across neurons.

A second, related possibility is that temporal pattern distinctiveness is an outcome of differences in cognition between events with respect to stimulus onset (e.g., onset of item A in the present study), and thus is not necessarily a product of a pattern separation computation per se. Lapses in attention prior to an event (deBettencourt et al., 2018; Kafkas and Montaldi, 2011; Kucewicz et al., 2018; Madore et al., 2020; Madore and Wagner, 2022; Miller and Unsworth, 2020), differences in what or where attention is allocated to during an event (Aly and Turk-Browne, 2016; deBettencourt et al., 2021; Dudukovic et al., 2011; Günseli and Aly, 2020; Olypher et al., 2002), and changes in motivational state or goals (Amer and Davachi, 2023; Kennedy and Shapiro, 2009) could impact the timing of cortical input into the hippocampus and therefore elicit more distinct temporal patterns across events. How to differentiate between effects of variability in cognition and stimulus processing over time and effects of a pattern separation computation on temporal pattern distinctiveness in the hippocampus remains an important objective for future research.

A third possibility is that changes in temporal pattern distinctiveness emerge from changes in single cell firing rates. Data from ex-vivo rodent hippocampal slices suggest that changes in firing rate in support of pattern separation may be more prevalent at longer timescales relative to pattern separation supported by spike timing reorganization (Madar et al., 2019a, 2019b). Studies of rate remapping in rodents indicate that hippocampal cells can change their firing rates in response to changes in the environment (Colgin et al., 2008). Even in the restricted case where the same set of cells are firing in the same order, changes in single cell firing rates can cause the population level temporal patterns of activity to change. In both cases of changes in firing sequence and firing rate, increased distinctiveness in temporal patterns in the hippocampus may be, in part, an outcome of extra-hippocampal brain regions exerting direct or indirect influence on hippocampal inputs or processes (Amer and Davachi, 2023).

A fourth explanation is that, even though wTPS was computed at the electrode level, changes in temporal patterns as observed from LFP recordings may nonetheless reflect changes in neural patterns of activity along a fine-scale spatial dimension. Single-cell recordings from humans indicate that hippocampal neurons can exhibit high selectivity in their responses to individual stimuli (Quiroga et al., 2005; Suthana et al., 2015). Findings from rodent studies document the flexibility in place field representations through global remapping, providing further evidence for the possibility of changes in neural ensembles in response to changes in the environment (Colgin et al., 2008). In the present case, the differential temporal patterns elicited by two different stimuli could be an emergent property of having different groups of neurons firing sequentially with their individual signatures at each timepoint; the degree of distinctiveness in the spatial patterns could thus be reflected in the distinctiveness in the temporal patterns. Furthermore, if we consider hippocampal activity as following attractor dynamics, for all potential mechanisms presented above, it is possible that the differential temporal patterns can be driven by either the presented stimuli, the starting positions of attractor dynamics, or both. While we offer multiple mutually compatible explanations for the presently observed changes in temporal patterns of neural activity as recorded from LFPs, the underlying neural computations governing the observed changes in hippocampal temporal pattern distinctiveness as it relates to subsequent memory behavior remain an open question.

Beyond this fundamental question, we note that subsequent memory was related to hippocampal temporal pattern distinctiveness during the encoding of the first item (A) in a sequential A-B event, whereas a null relationship was observed between item B pattern distinctiveness and subsequent memory. On the one hand, we might expect the hippocampal-dependent conjunctive representations that support later memory to form, at least in part, during the item B period. Indeed, results from extant fMRI studies using similar sequential paired-associate encoding tasks (but with longer lags between the A and B items than in the present case) suggest that hippocampal activity during item B encoding is associated with subsequent associative memory (Hales et al., 2009; Hales and Brewer, 2010; Qin et al., 2009; Staresina and Davachi, 2009). Extending these findings, we also observed that HFB power during B items predicted future memory behavior (Fig. 2B). On the other hand, the present absence of a relationship between item B wTPS and subsequent memory may reflect the present task structure, wherein additional cognitive operations were likely engaged during the item B period compared to the item A period. For example, it is likely that participants either maintained and/or retrieved the item A representation during the item B encoding period, resulting in a potential conflation of neural signals during the item B period. Consistent with this possibility, we note that the effects of stimulus category on HFB power were qualitatively different in the two encoding periods (see Fig. S6, and contrast with the qualitatively similar HFB category effects observed for the study phase item A period and the test phase cue-locked period). The additional processes engaged during the item B period likely interfered with our ability to detect and isolate effects of temporal pattern similarity during item B encoding.

Another fundamental question is whether hippocampal temporal pattern distinctiveness differentially predicts future pattern completion-dependent (e.g., associative hits/cued recall) versus familiarity-dependent (e.g., item recognition) memory behaviors. Data from some prior human fMRI studies show that hippocampal activity at encoding differentially predicts later conjunctive memory expression (e.g., source recollection) relative to item recognition (Davachi et al., 2003; Kirwan and Stark, 2004; Ranganath et al., 2004), with other studies showing that subsequent memory strength scales with hippocampal activity (Shrager et al., 2008; Song et al., 2011). Debate remains about how to map behavioral expressions of memory to specific computations (e.g., pattern completion vs. summed-similarity familiarity computations). The present results further link HFB power at both encoding and retrieval to memory behavior. Using a two-step old/new recognition-cued recall retrieval paradigm, we observed a graded increase in the positive HFB power subsequent memory effect at encoding (associative hits > item hits > misses), and a difference in retrieval HFB power between associative hits and item hits, with item hits and CRs not differing. Complementing these univariate effects, we show that hippocampal temporal distinctiveness during the item A period differentiated subsequent associative hits and item hits from misses. Interestingly, the temporal similarity during the item A period did not differentiate associative hits and item hits in the same way as HFB power, suggesting that there may be potential differences in the timescales for how HFB power and temporal pattern similarity represent information encoding that will need to be explored further in future studies. Given the recognition-cued recall nature of the paradigm along with our coding of the behavioral responses, item hits, while not accompanied by pattern completion of specifics about the associate (B item), were based on a mixture of pattern-completion dependent (e.g., retrieval of other event features) and summed-similarity familiarity computations. Future studies that assay familiarity-based item recognition, controlling for pattern-completion dependent memory, hold promise for determining whether hippocampal pattern separation at encoding selectively fosters future pattern completion or also impacts future familiarity.

Extant fMRI studies examining content representations in the human hippocampus have produced variable results (Bonnici et al., 2012; Diana et al., 2008; LaRocque et al., 2013; Liang et al., 2013; Rissman and Wagner, 2012). While some studies have shown that content at the category level (e.g., scenes versus faces) could not be decoded from distributed BOLD patterns in the hippocampus (Diana et al., 2008; LaRocque et al., 2013; Liang et al., 2013), Bonnici and colleagues (2012) found distinct hippocampal representations for complex scenes being viewed. The present exploratory analyses revealed effects of stimulus category on univariate HFB power at both encoding and retrieval. While informative, it is important to note that these results are from group-level analyses using mixed-effects models over random participant-channel effects; the electrode coverage of different regions of the hippocampus across participants was insufficient to enable examination of category effects as a function of anatomical location. Future studies are needed to further clarify the relationship between hippocampal subregions (e.g. along the rostrocaudal axis) and stimulus category induced HFB power. Furthermore, while the present study is focused solely on the temporal dimension of pattern similarity, the relationship between hippocampal processes in the temporal and spatial dimensions warrant further investigation.

Taken together, our findings add to a rich literature focused on the computations of human hippocampus that support memory, a literature to which Eleanor Maguire made numerous significant contributions. From characterization of hippocampal mismatch (prediction error) computations (Garrido et al., 2015; Kumaran and Maguire, 2007, 2006) to pattern separation (Chadwick et al., 2011) to pattern completion (Bonnici et al., 2012), Eleanor was at the forefront of leveraging human functional imaging to test models of hippocampal computation. Moreover, Eleanor was a developer and early adopter of emerging tools and approaches, including advocating for the field to shift to more naturalistic paradigms (Maguire, 2012) and demonstrating the power of multivariate analyses, which led to many new insights, including that patterns of hippocampal BOLD activity differ across events and thus are decodable (Chadwick et al., 2012, 2010). Her field-leading work provided a foundation for future studies of hippocampal replay (e.g., Tambini and Davachi, 2019) and prospective coding (e.g., Brown et al., 2016), amongst others. Eleanor’s creative science and novel discoveries inspired and informed our science (e.g., Chen et al., 2015, 2011; Xue et al., 2026) and undoubtedly that of many others. Indeed, the present study complements Eleanor’s exploration of how mnemonic representations are encoded in hippocampal spatial patterns, documenting how later remembering also is associated with hippocampal pattern distinctiveness that operates along the temporal dimension.

## Data and Code Availability

Bayesian model specifications and outputs are accessible at https://github.com/alicexue/temporal-encoding-patterns. The data and preprocessing code will be archived and made available via the Stanford Digital Repository upon request.

## Author Contributions

Alice M. Xue: Formal Analysis, Visualization, Writing – Original Draft Preparation

Shaw Hsu: Formal Analysis, Project Administration, Software, Visualization, Writing – Original Draft Preparation

Karen F. LaRocque: Conceptualization, Formal Analysis, Methodology, Software, Writing – Review & Editing Omri Raccah: Investigation, Project Administration, Methodology, Writing – Review & Editing

Alex Gonzalez: Conceptualization, Investigation, Methodology, Software, Writing – Review & Editing

Josef Parvizi: Conceptualization, Formal Analysis, Funding Acquisition, Methodology, Supervision, Writing – Review & Editing

Anthony D. Wagner: Conceptualization, Formal Analysis, Funding Acquisition, Methodology, Supervision, Writing – Original Draft Preparation

**The authors declare no conflict of interest.**

## Acknowledgments

Supported by grants from the National Institute of Neurological Disorders & Stroke (R01-NS078396), National Science Foundation (BCS-1358907), Marcus and Amalia Wallenberg Foundation, National Science Foundation Graduate Fellowships (to A.M.X., K.F.L., O.R., & A.G.), Stanford Interdisciplinary Graduate Fellowships (to A.M.X. & S.H.), Stanford DARE Fellowship (to A.G.), and the Stanford Center for Mind, Brain, and Computation (to A.M.X., S.H., K.F.L., & A.G.). We thank Janice Chen, Jintao Sheng, Alvin Tan, and Tobi Gerstenberg for helpful discussions, the Stanford Epilepsy Monitoring Unit Staff for assistance with data collection, and the patients for donating their time during clinical evaluation at the Stanford Medical Center.

## Supplementary Material

### Effects of Stimulus Category on Memory Performance

The studied paired associates consisted of items drawn from four categories — famous faces, famous buildings, numbers, and words — with the A and B terms drawn from different categories (see *Material and Methods*). Using a series of linear mixed-effects models predicting d’ and associative hit rate/cued recall probability as a function of various item category predictors (see *Material and Methods* for models and control variables), we first observed that inclusion of item A category in a model predicting memory performance improved predictive performance for d’ (Fig. S4A; ELPD = −29.52, ΔELPD = −14.86 ± 4.10) and associative hit rate (Fig. S4B; ELPD = 16.46, ΔELPD = −5.70 ± 3.84). Estimated marginal contrasts revealed that d’ for number cues was lower than d’ for faces, buildings, and words (Δ = −2.337 [−3.031, −1.683]; Δ = −1.895 [−2.537, −1.228]; Δ = −1.541, [−2.238, −0.877]); associative hit rate when cued by number also was lower than associative hit rate cued by faces, buildings, and words (Δ = −0.122 [−0.245, −0.003]; Δ = −0.164, [−0.272, −0.039]; Δ = −0.258 [−0.377, −0.132]. These results indicate that recognition memory and cued recall suffered when the item A category was numbers. In addition, d’ for word cues was lower than d’ for faces (Δ = −0.793 [−1.410, −0.078]) and associative hit rate when cued by word was higher than associative hit rate cued by faces (Δ = 0.132 [0.010, 0.248]). At the same time, we note that the associative hit rate as a function of item A category was impacted by recognition memory of item A. A follow-up linear mixed effects model that excluded ‘miss’ trials revealed that associative hit rate when cued with words was greater than that when cued with faces, buildings, and numbers (Fig. S4C; Δ = 0.222 [0.100, 0.348]; Δ = 0.174 [0.052, 0.295]; Δ = 0.313 [0.198, 0.445]) and associative hit rate when cued with buildings was greater than that when cued with numbers (Δ = 0.138 [0.025, 0.259]).

Finally, we used a linear mixed-effects model to examine whether associative hit rate varied as a function of item B category. Model comparison revealed that the item B category predictor did not meaningfully improve predictive performance (Fig. S5A; ELPD = 28.25, ΔELPD = −2.10 ± 1.98). A follow-up mixed-effects model that excluded ‘miss’ trials — thus controlling for recognition memory for item A — showed that an item B category predictor marginally improved predictive performance on cued recall (Fig. S5B; ELPD = 18.92, ΔELPD = −0.89 ± 2.78).

### Effects of Stimulus Category on Univariate Hippocampal HFB Power

We examined the effects of both item A and item B categories on univariate HFB power. Estimated marginal contrasts between categories and HFB time courses are shown in S6, S7. For the study phase, inclusion of an item A category predictor substantially improved predictive performance for the item A period (ELPD = −19332.07, ΔELPD = −71.14 ± 12.31). Estimated marginal contrasts showed that HFB power for words was lower than for faces and buildings (Δ = −0.046 [−0.055, −0.037]; Δ = −0.042 [−0.051, −0.033]); HFB power for numbers was lower than for faces and buildings (Δ = −0.038 [−0.048, −0.029]; Δ = −0.034, [−0.043, −0.024]). There was also a substantial improvement in predictive performance when an item B category predictor was included in a model of the item B period (ELPD = −21095.94, ΔELPD = −43.95 ± 9.90). Estimated marginal contrasts showed that HFB power for numbers was higher than for faces, buildings, and words (Δ = 0.034 [0.024, 0.044]; Δ = 0.036 [0.027, 0.045]; Δ = 0.045 [0.036, 0.055]); HFB power for words was lower than for faces (Δ = −0.011 [−0.022, −0.003]).

For the test phase, predictive performance for the cue-locked period was substantially improved when item A and item B category predictors were included (ELPD = −9529.73; ΔELPD for item A = −57.20 ± 10.87; ΔELPD for item B = −31.16 ± 8.30). Estimated marginal contrasts for item A category showed that HFB power for buildings was higher than for numbers and words (Δ = 0.057 [0.045, 0.069]; Δ = 0.045 [0.033, 0.056]), but not faces (Δ = 0.008 [−0.003, 0.020]); HFB power for faces was higher than for numbers and words (Δ = 0.049 [0.036, 0.060]; Δ = 0.037, [0.026, 0.048]). Estimated marginal contrasts for item B category showed that HFB power for words was lower than for faces, buildings, and numbers (Δ = −0.046 [−0.056, −0.034]; Δ = −0.016 [−0.028, −0.005]; Δ = −0.032, [−0.044, −0.021]); HFB power for buildings was less than faces and numbers (Δ = −0.029 [−0.041, −0.018]; Δ = −0.016 [−0.027, −0.004]). Model comparison of the response-locked period showed that predictive performance improved with the inclusion of item A and item B category predictors (ELPD = −11971.92; ΔELPD for item A = −3.63 ± 3.65; ELPD for item B = ΔELPD = −12.95 ± 5.67). Estimated marginal contrasts for item A category showed that HFB power for numbers was lower than for faces, buildings, and words (Δ = −0.017 [−0.030, −0.003]; Δ = −0.023 [−0.036, −0.009]; Δ = −0.021 [−0.035, −0.008]). Estimated marginal contrasts for item B category showed that HFB power for faces was higher than for buildings and words (Δ = 0.027 [0.015, 0.040]; Δ = 0.031 [0.018, 0.044]); HFB power for numbers was higher than for buildings and words (Δ = 0.018 [0.005, 0.030]; Δ = 0.022 [0.009, 0.034]).

Collectively, these data reveal that during item A encoding and during retrieval cuing with item A at test, human hippocampal HFB power was greater for buildings (famous landmarks) and faces (of famous individuals) than for words and numbers. The more complex effects of item B stimulus category at encoding and at test likely reflect (1) the blending of item B encoding and the maintenance and/or reinstatement of item A (at study) and (2) the blending of item A encoding and the reinstatement of item B (at retrieval).

### Effect of Participant S5 on Hippocampal HFB Temporal Pattern Similarity and Subsequent Memory Results

Participant S5 was included in all temporal pattern similarity linear mixed-effects models presented in the results section, even though some datapoints from that participant are clearly outliers. To explore whether the critical findings of lower wTPS during the encoding of subsequent associative hits and item hits relative to subsequent misses hold, excluding S5, we recomputed these analyses. The model with item A wTPS as the outcome and subsequent memory as a predictor (ELPD = −3323.18) outperformed the model without subsequent memory, though this difference was small (ΔELPD = −3.34 ± 3.17). Estimated marginal contrasts showed that item A wTPS for subsequent misses was greater than that for subsequent associative hits and item hits (Δ = 0.140 [0.013, 0.258]; Δ = 0.178 [0.071, 0.283]), which did not differ from each other (associative hit – item hit = 0.039 [−0.055, 0.126]). Similarly, the linear mixed-effects model for item A aTPS was re-run with participant S5 excluded and the results did not qualitatively change; inclusion of subsequent memory outcome as a predictor did not improve predictive performance (ELPD for the model without subsequent memory = −3321.52, ΔELPD = −1.78 ± 0.82). Additional models showed that the similarity type (wTPS vs aTPS) × memory outcome interaction did not improve predictive performance (ELPD for the model without the interaction term = −6644.84, ΔELPD = −0.18 ± 1.99) for the associative hit and miss contrasts (Δ = −0.091 [−0.258, 0.059]) and item hit and miss contrasts (Δ = −0.138 [−0.273, 0.003]). Taken together, these analyses show that the results hold when data from participant S5 were excluded.

### Effect of Participants S2 and S5 on Hippocampal HFB Temporal Pattern Similarity and Subsequent Memory Results

Participants S2 and S5 both showed lower associative hit rates compared to the other participants. While their recognition d’ and CVLT delayed recall retention scores suggest that their low associative hit rates may be due to high decision thresholds instead of poor associative memory (see Results in Main Text), there remains the possibility that these two participants’ data are unrepresentative of the population for this memory task. To explore whether the critical findings of lower wTPS during the encoding of subsequent associative hits and item hits relative to subsequent misses is observed independent of their data, we recomputed these analyses excluding S2 and S5. The model with item A wTPS as the outcome showed better performance with subsequent memory as a predictor (ELPD = −2990.50) than a model without subsequent memory as a predictor, though again, this difference was small (ΔELPD = −2.65 ± 2.92). Estimated marginal contrasts showed that item A wTPS for subsequent misses was greater than that for subsequent associative hits and item hits (Δ = 0.148 [0.015, 0.279]; Δ = 0.182 [0.072, 0.309]), which did not differ from each other (associative hit – item hit = 0.035 [−0.053, 0.130]). Similarly, the linear mixed-effects model for item A aTPS was recomputed with participants S2 and S5 excluded and the results did not qualitatively change; inclusion of subsequent memory as a predictor did not improve predictive performance (ELPD for the model without subsequent memory = −2984.22, ΔELPD = −1.52 ± 0.90). Estimated marginal contrasts showed that the similarity type (wTPS vs aTPS) × memory outcome interaction did not improve predictions of similarity (ELPD for the model without the interaction = −5975.17, ΔELPD = −0.69 ± 1.70) for the associative hit and miss contrasts (Δ = −0.091 [−0.260, 0.081]) and item hit and miss contrasts (Δ = −0.130 [−0.281, 0.017]). Taken together, these analyses show that the results hold when data from participants S2 and S5 were excluded.

### Effect of Alternative Categorization of Memory Outcomes on HFB Memory Analyses

To explore how the categorization of memory outcomes may have impacted the results, we recomputed the critical univariate and multivariate memory analyses by grouping memory response categories (2) and (3) into the “item hit” condition and response categories (4) and (5) into the “associative hit” condition (see *Materials and Methods* for definitions and original memory outcome categorization).

First, the univariate hippocampal HFB analyses were recomputed. For the study phase (Fig. S10A, B), inclusion of a subsequent memory predictor improved predictive performance for both the item A (ELPD = −19332.07, ΔELPD = −18.67 ± 6.36) and item B (ELPD = −21088.43, ΔELPD = −25.60 ± 7.36) encoding periods. Estimated marginal contrasts showed that (a) HFB power for subsequent associative hits was higher than both subsequent items hits and misses during encoding of item A (Δ = 0.009 [0.001, 0.016]; Δ = 0.033 [0.023, 0.043]) and during encoding of item B (Δ = 0.019 [0.011, 0.026]; Δ = 0.040 [0.030, 0.051]); and (b) HFB power for item hits was higher than misses during encoding of item A (Δ = 0.025 [0.017, 0.035]) and during encoding of Item B (Δ = 0.021 [0.012, 0.031]). For the test phase (Fig. S10C, D), a memory (i.e., retrieval success) predictor for both cue-locked and response-locked periods improved predictive performance (ELPD = −15784.84, ΔELPD = −11.61 ± 5.54; ELPD = −19908.32, ΔELPD = −105.27 ± 14.68). Estimated marginal means showed that (a) HFB power for associative hits was higher than both item hits and CRs during the cue-locked (Δ = 0.015 [0.007, 0.024; Δ = 0.022 [0.013, 0.030]) and response-locked (Δ = 0.066 [0.057, 0.076]; Δ = 0.057 [0.047, 0.066]) periods; and (b) HFB power for item hits was higher than CR during the response-locked period (Δ = −0.009 [−0.017, −0.001]), but not during the cue-locked period (Δ = 0.007 [−0.001, 0.014]).

The analyses on the relationships between temporal pattern similarity and subsequent memory were also recomputed (Fig. S11). The model with item A wTPS as the outcome and subsequent memory outcome as a predictor (ELPD = −3405.47) modestly outperformed a model without memory outcome as a predictor (ΔELPD = −4.36 ± 3.49). Estimated marginal means showed that item A wTPS for subsequent misses was greater than for item hits (Δ = 0.186 [0.084, 0.292]) but not associative hits (Δ = 0.110 [−0.011, 0.234]); wTPS for subsequent item hits and associative hits did not differ from each other (associative hit – item hit = 0.076 [−0.013, 0.164]). Similarly, the linear mixed-effects model for item A aTPS was re-run; subsequent memory outcome as a predictor did not improve predictive performance (ELPD = −3414.49, ΔELPD = −1.43 ± 0.89). Additional models showed that the similarity type (wTPS vs aTPS) × memory outcome interaction did not improve predictive performance (ELPD = −6819.14, ΔELPD = −0.41 ± 1.77) for the associative hit and miss contrast (Δ = −0.099 [−0.254, 0.047]) nor for item hit and miss contrast (Δ = −0.125 [−0.271, 0.009]).

### Effect of Aggregating Associative Hit and Item Hit Categories on Temporal Pattern Similarity Analyses

To alleviate any potential concern that the pattern similarity analyses were underpowered due to low trial counts for associative hits and/or item hits and to ensure that the results presented here are not driven by the specific memory outcome categorizations used, we recomputed these analyses after aggregating associative hit and item hit categories into a single ‘hit’ category (Fig. S12). The model with item A wTPS as the outcome and subsequent memory as a predictor revealed a modest improvement in predictive performance over a model without subsequent memory (ELPD = −3406.00, ΔELPD = −3.92 ± 3.03). Similarly, the linear mixed-effects model for item A aTPS was recomputed with only the ‘miss’ and aggregated ‘hit’ memory categories and the results did not qualitatively change; that is, subsequent memory outcome included as a predictor did not improve predictive performance (ELPD = −3414.49, ΔELPD = −0.84 ± 0.33). Additional models showed that the similarity type (wTPS vs aTPS) × memory outcome interaction did not improve a model of similarity (ELPD = −6819.14, ΔELPD = −0.41 ± 1.77) for the associative hit and miss contrast (Δ = −0.099 [−0.254, 0.047]) and item hit and miss contrast (Δ = −0.125 [−0.271, 0.009]). Taken together, these analyses show that the pattern similarity results hold when we maximized statistical power by aggregating associative hits and item hits, though effects were more modest, likely due to the aggregation of memories of varying strength in the ‘hit’ condition.

## Supplemental Figures

**Figure S1.**
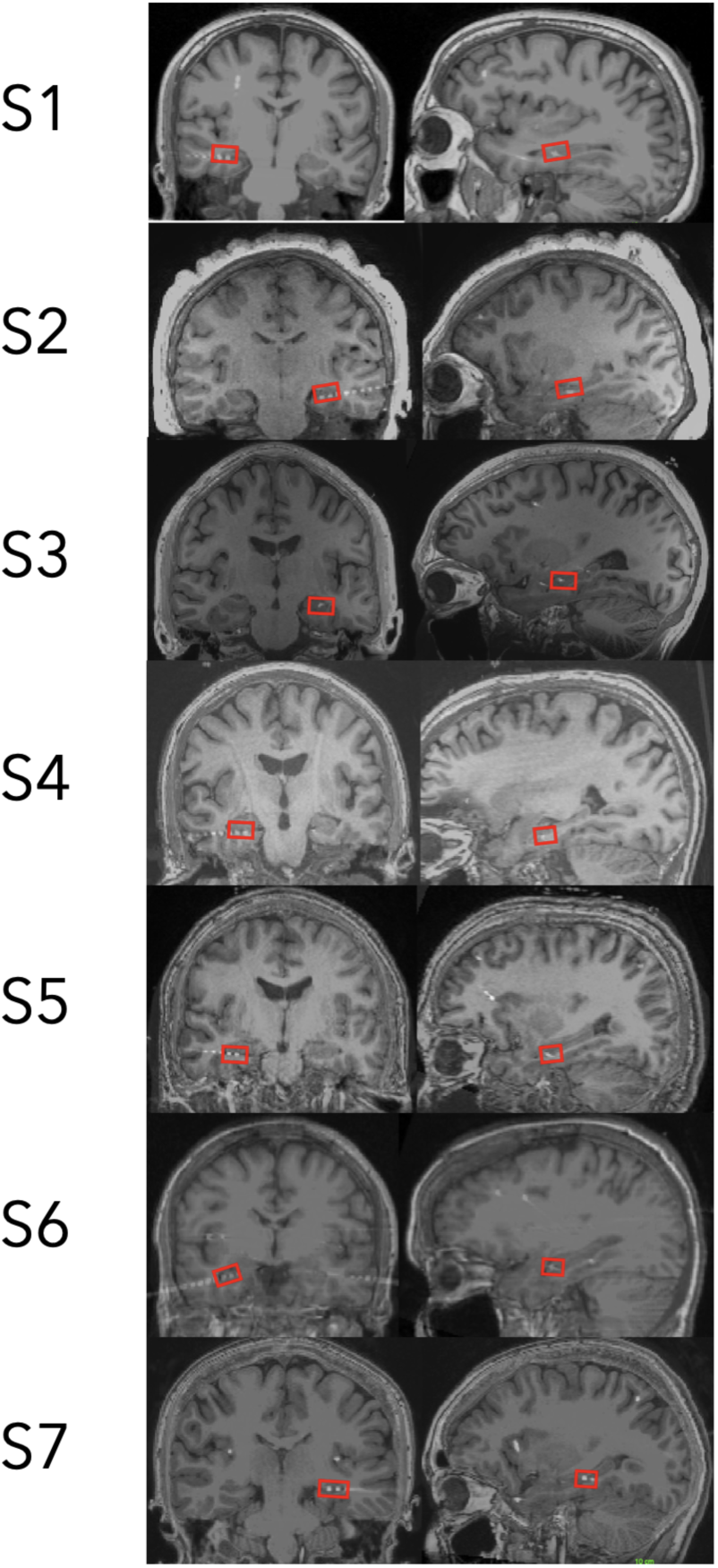
Example hippocampal electrodes in each participant. Electrodes were visualized using the ITK-SNAP software. Electrode localization and selection were guided by overlaying post-surgical CT images of electrodes that were aligned with structural MRI of the brain for each participant.

**Figure S2.**
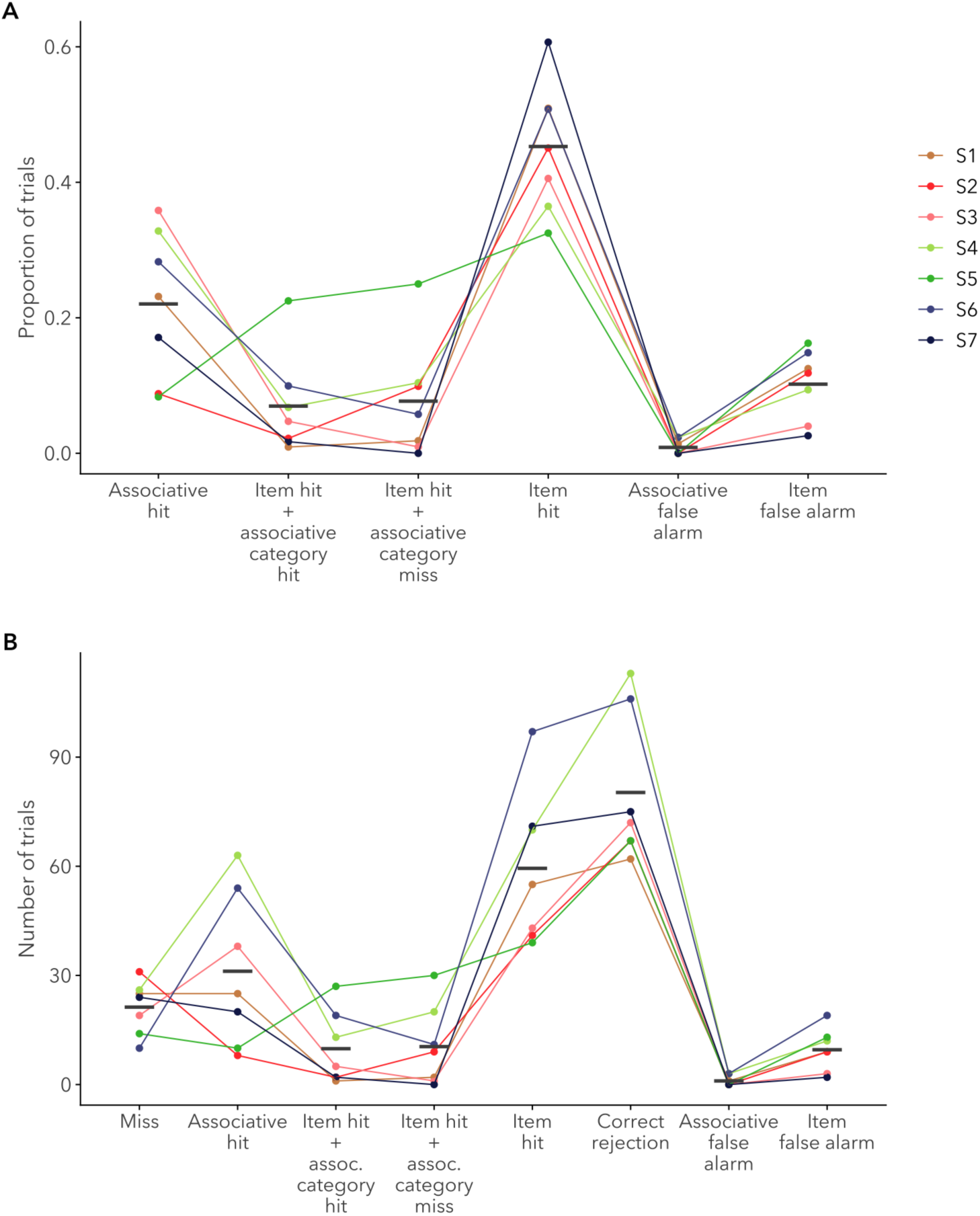
Behavioral results prior to aggregation as both (A) rates and (B) trial counts. Each point denotes an individual participant and line segments denote weighted group means. Associative hit: responded to a studied cue with “old, remember something about its pair,” correctly identified the category of the pair, and correctly generated at least one qualitative description of the associate during verbal probe; Item Hit + Associative Category Hit: responded to a studied cue with “old, remember something about its pair” and correctly identified the category of the pair but was unable to correctly generate at least one qualitative description of the associate during verbal probe; Item Hit + Associative Category Miss: responded to a studied cue with “old, remember something about its pair” but failed to correctly identify the category of the pair during verbal probe; Item Hit: responded to a studied cue with “old, remember nothing about its pair”; Associative False Alarm: responded to a novel cue with “old, remember something about its pair”; Item False Alarm: responded to a novel cue with “old, remember nothing about its pair.”

**Figure S3.**
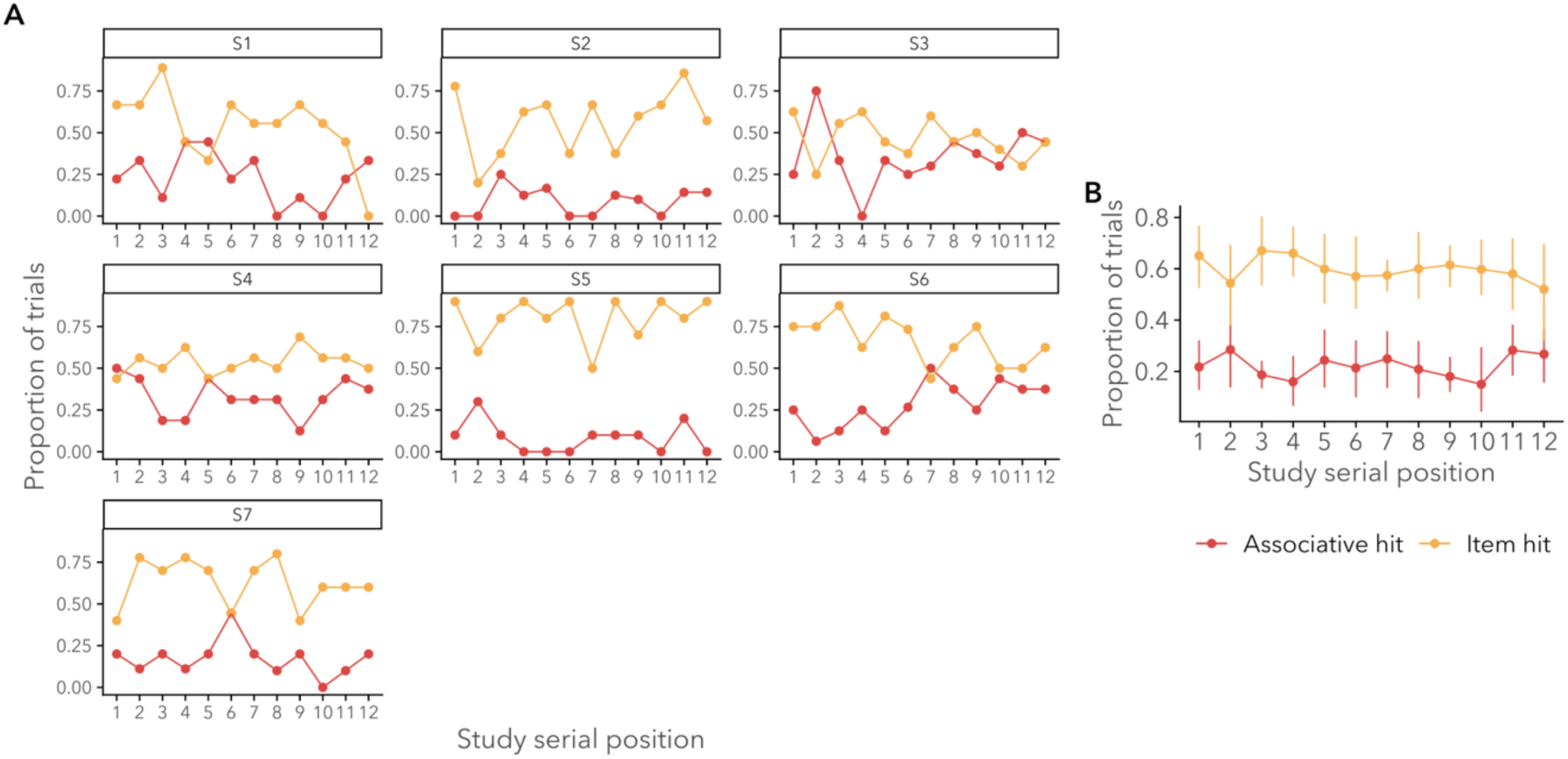
Proportion of trials for each behavioral outcome as a function of study serial position. (A) Serial position effects for individual participants. (B) Mean serial position effects across participants. Error bars represent bootstrapped 95% confidence intervals.

**Figure S4.**
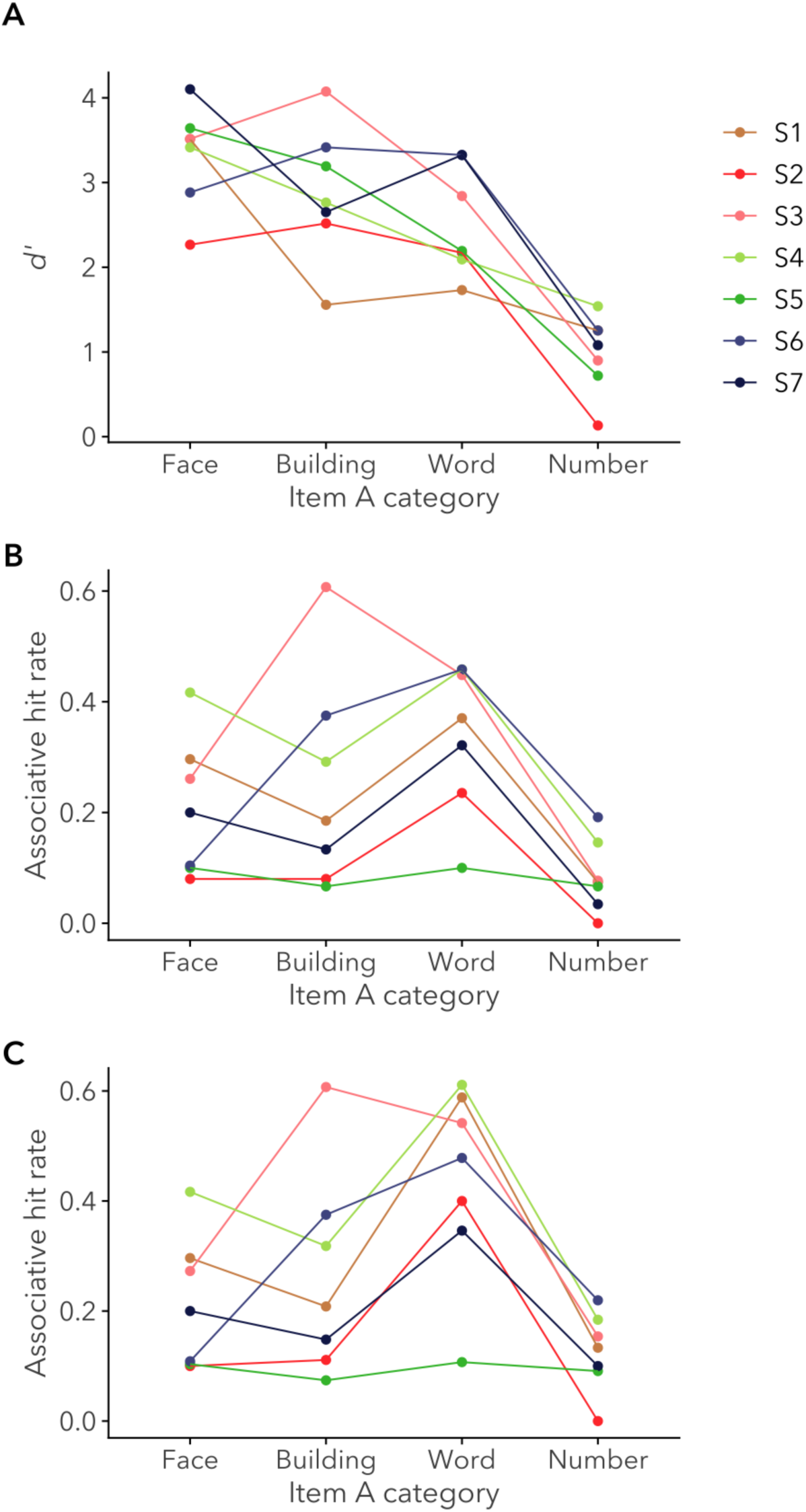
Memory performance as a function of item A category: (A) d’; (B) Associative hit rate; (C) Associative hit rate for correct ‘old’ trials. Each point denotes an individual participant.

**Figure S5.**
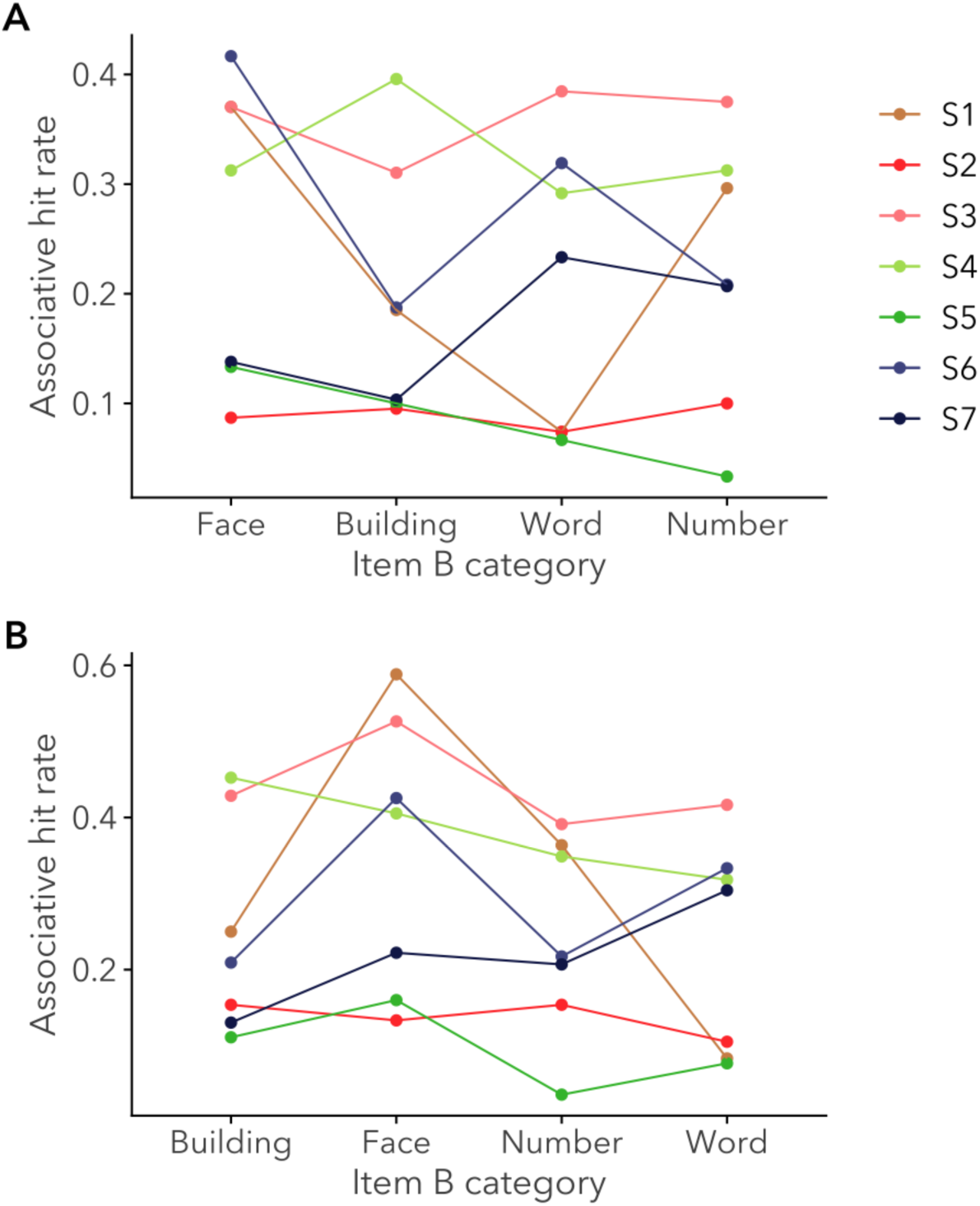
Memory performance as a function of item B category: (A) Associative hit rate; (B) Associative hit rate for correct ‘old’ trials. Each point denotes an individual participant.

**Figure S6.**
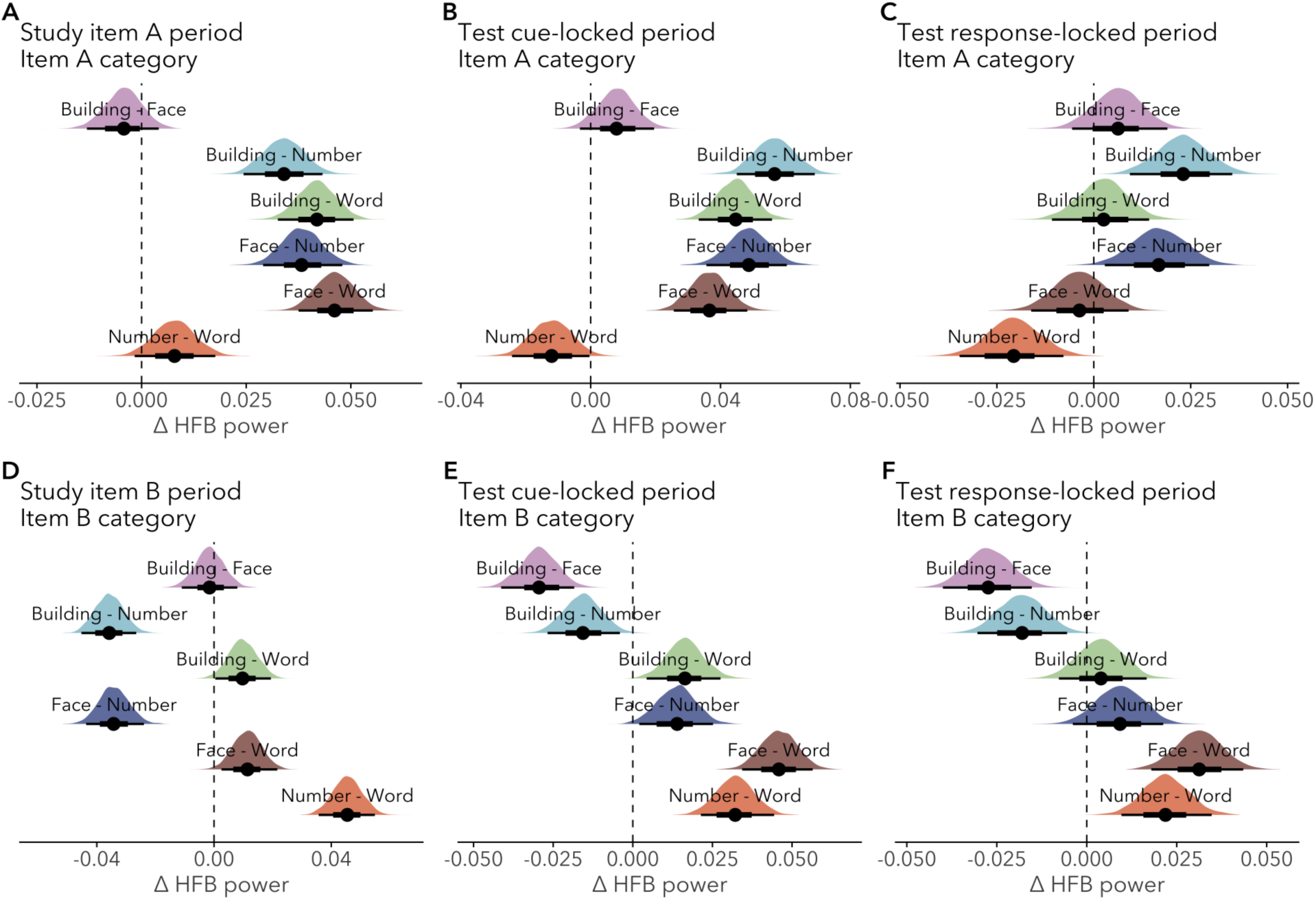
Estimated marginal contrasts for univariate HFB between stimulus categories for study and test phases. Posterior probability distributions for estimated marginal contrasts are depicted in the shaded areas; points indicate posterior medians and horizontal lines indicate the 66% and 95% HDIs.

**Figure S7.**
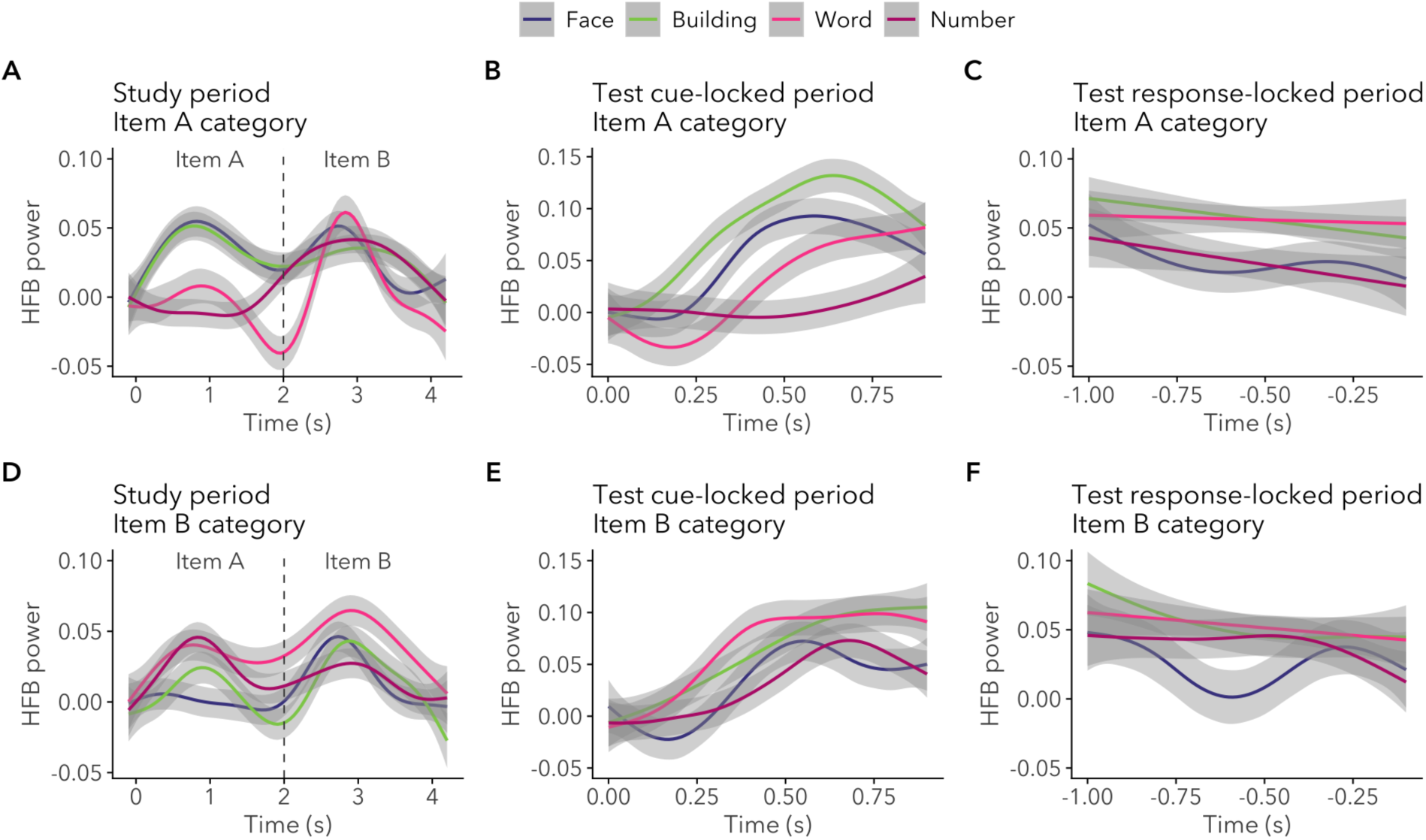
Timeseries plot of HFB power for each stimulus category for study and test phases. Lines are smoothed for visualization purposes. Error bars represent bootstrapped 95% confidence intervals.

**Figure S8.**
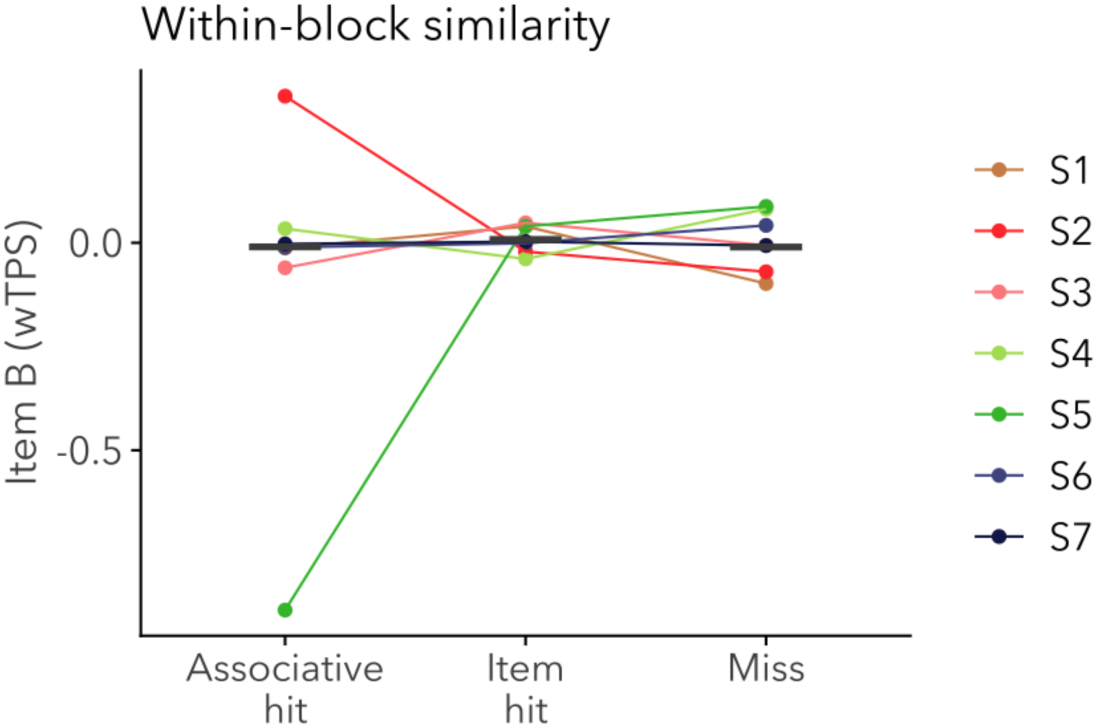
Within-block temporal pattern similarity (wTPS) for item B as a function of subsequent memory. Each point denotes an individual participant and line segments denote weighted group means.

**Figure S9.**
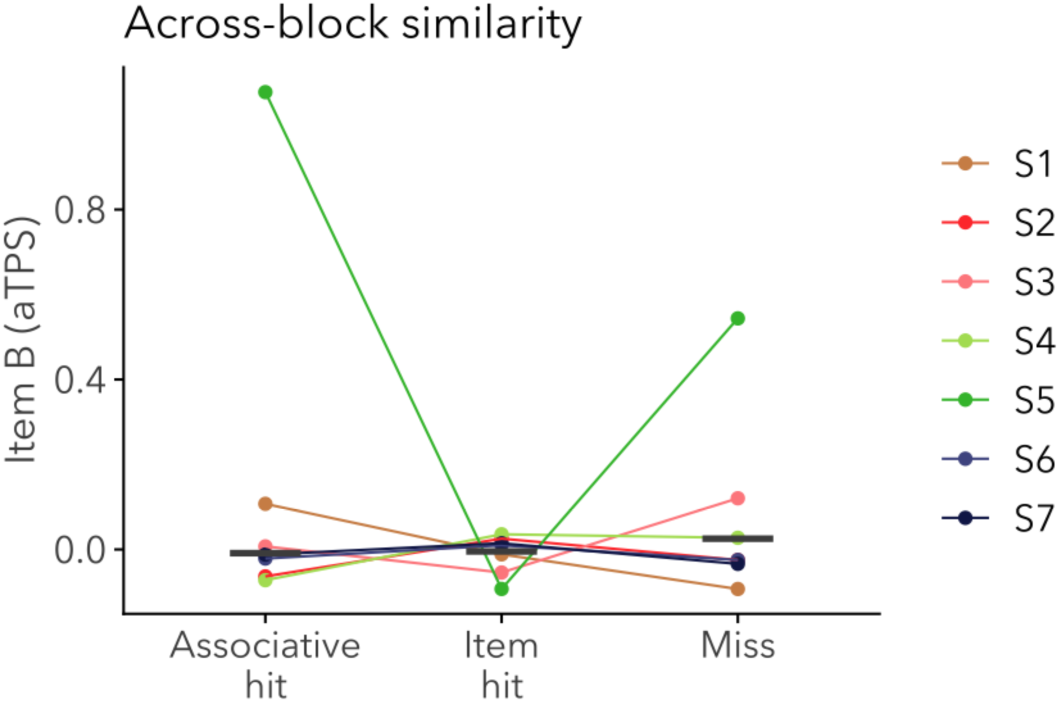
Across-block temporal pattern similarity (aTPS) for item A as a function of subsequent memory with axis scaled to show outlier data points from participant S5. Each point denotes an individual participant and line segments denote weighted group means.

**Figure S10.**
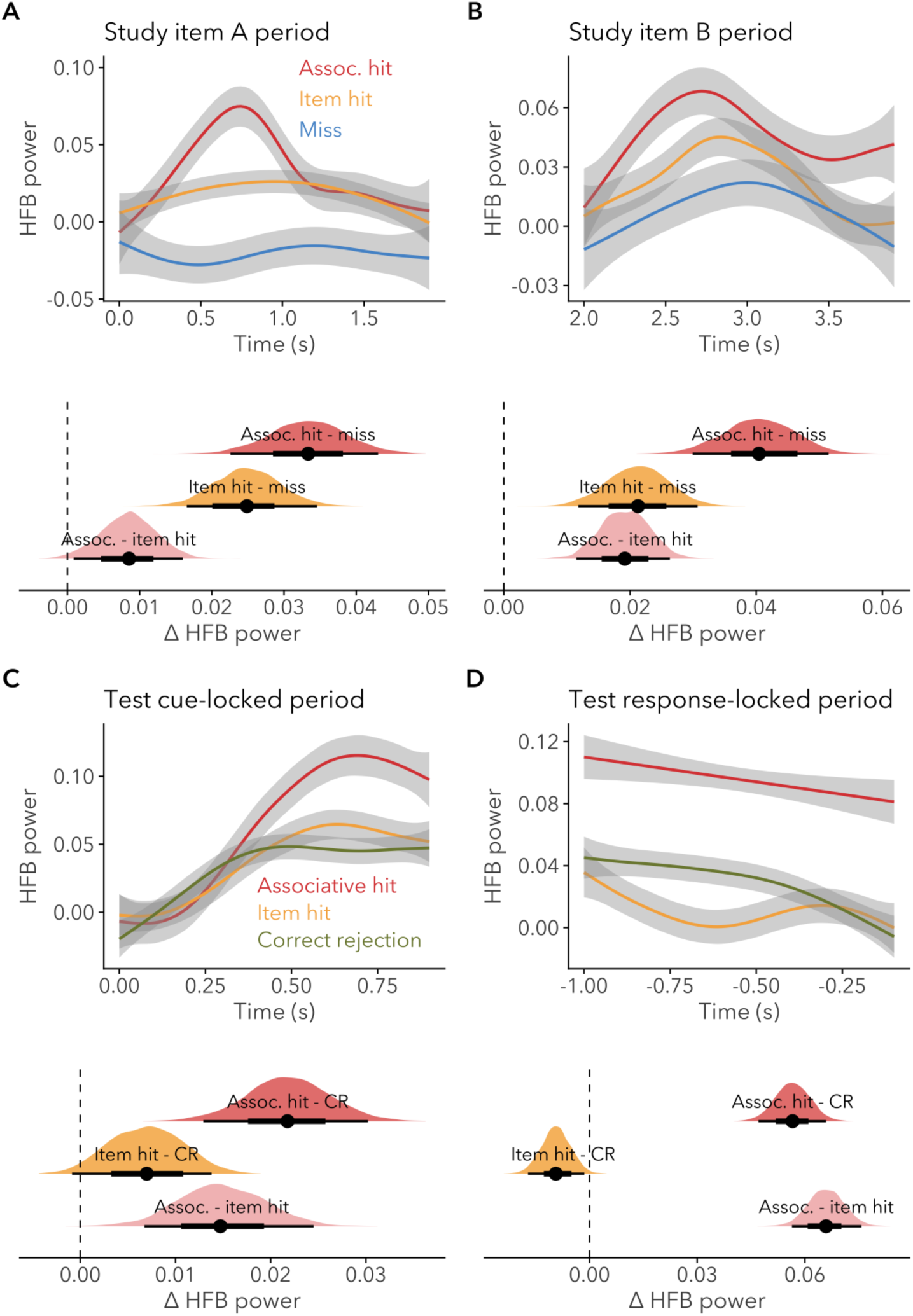
Univariate HFB timeseries plots (top) and estimated marginal contrasts for HFB (bottom) between alternative memory outcomes during the (A, B) study phase and (C, D) test phase. Timeseries were smoothed for visualization purposes. Error bars on the timeseries represent bootstrapped 95% confidence intervals. Plots of estimated marginal contrasts show posterior probability distributions in the shaded areas; points indicate posterior medians and horizontal lines indicate the 66% and 95% HDIs.

**Figure S11.**
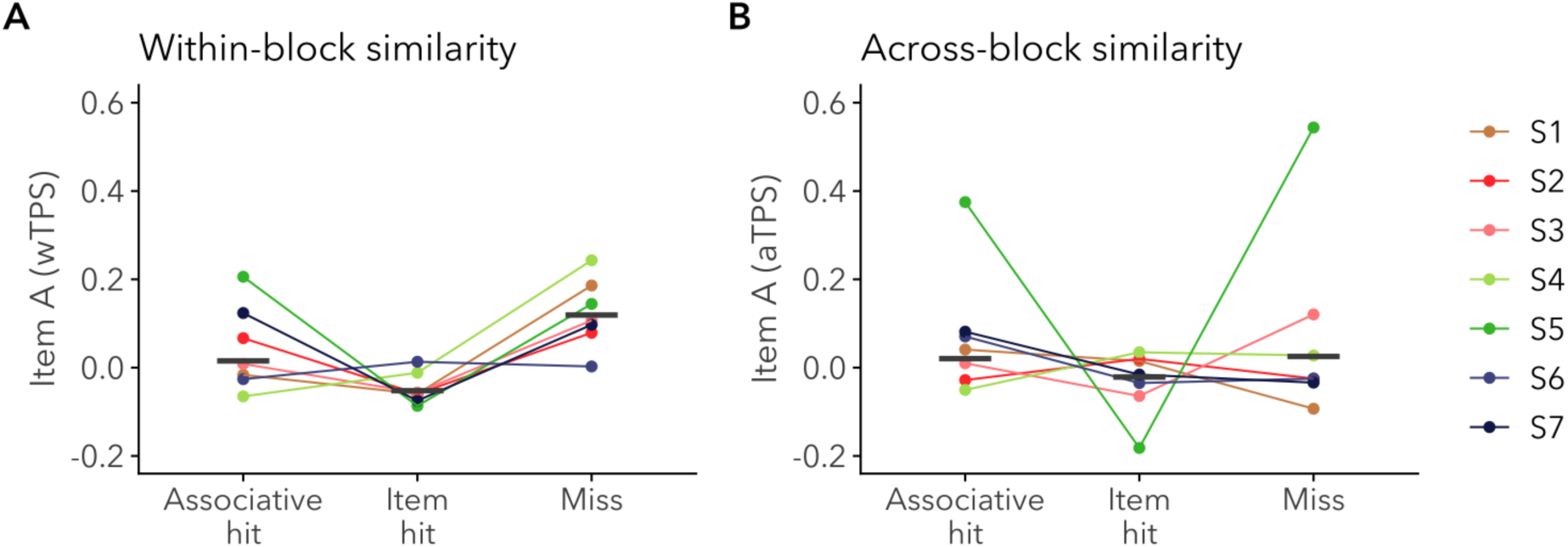
Study phase item A (A) within-block temporal pattern similarity (wTPS) and (B) across-block temporal pattern similarity (aTPS) with alternative memory outcome categorization. Each point denotes an individual participant and line segments denote weighted group means.

**Figure S12.**
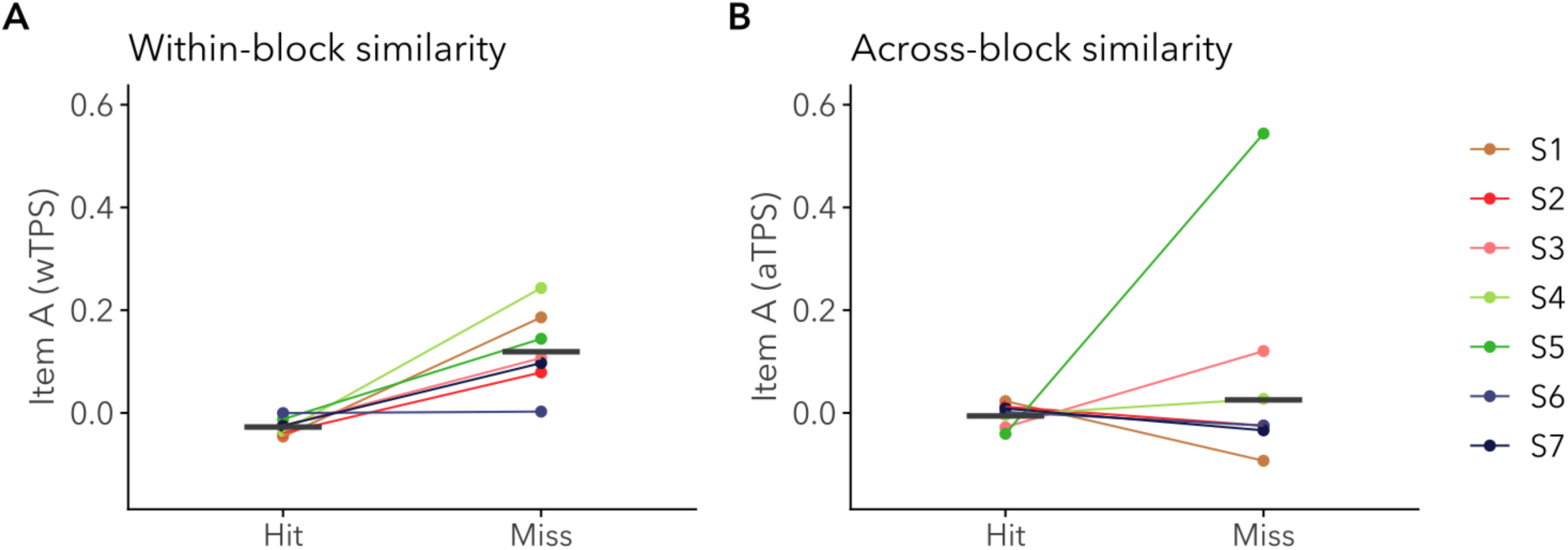
Study phase item A (A) within-block temporal pattern similarity (wTPS) and (B) across-block temporal pattern similarity (aTPS) with associative hits and item hits combined into a single hit category. Each point denotes an individual participant and line segments denote weighted group means.

